# The importance of visual and chemical cues in infection detection and avoidance in a freshwater fish

**DOI:** 10.1101/2024.03.27.587060

**Authors:** Ariane Côté, Sandra A. Binning

## Abstract

The characteristics of infected animals, including their smell, appearance, behaviour or sound, can greatly differ from those of uninfected conspecifics. These differences can serve as cues to recognize and avoid infected individuals to minimize the risk of infection. Avoidance of infected conspecifics is a risk-sensitive behaviour that can be influenced by various factors such as sensory cues and environmental parasite load, which both remain poorly understood in fish. We investigated the ability of two populations of wild caught pumpkinseed sunfish (*Lepomis gibbosus*) to distinguish between conspecifics infected with parasitic worms versus uninfected individuals in two-choice experiments using visual and chemical cues separately. One population was from a lake without parasites (parasite naive) whereas the other population originated from a lake with a high prevalence of trematode and cestode worms (parasite experienced). Both populations preferred to be with conspecifics, regardless of their infection level, over being alone when given visual cues but avoided conspecifics and remained alone when given chemical cues, suggesting that visual and chemical cues are not redundant. Neither population showed any preference between infected and uninfected conspecifics when given visual cues. However, in the presence of chemical cues, there was a great interindividual variation: some fish preferred uninfected conspecifics while others preferred infected ones. On average, naive fish avoided infected conspecifics whereas experienced fish did not show any preferences, suggesting that fish from lakes with high prevalence of infection habituate to infection cues. We suggest that pumpkinseeds use chemical rather than visual cues to discriminate between infected and uninfected conspecifics and make a shoaling decision. Our study highlights the importance of considering different sensory cues as well as parasite load when studying avoidance and shoaling behaviours, especially in a time of modifying sensory landscape and parasites abundance of freshwater ecosystems through global changes.

## Introduction

Parasites are ubiquitous organisms that live on or in a host and exploit its resources (Lewin, 1982). By doing so, they can inflict an energetic burden and dramatically affect host performance traits (Chrétien et al., 2022), including movement behaviours (Binning et al., 2017) and sociality (Barber & Dingemanse, 2010; Dubois & Binning, 2022), that may ultimately affect reproduction and survival (Marcogliese, 2004). As a result, there is pressure on social hosts to avoid parasitic infection. Infected animals can exhibit distinctive sensory cues, such as odd smell, appearance, behaviour and/or sound, facilitating recognition and avoidance by conspecifics to reduce infection risks, i.e. infection avoidance behaviour (Lopes et al., 2022; Stockmaier et al., 2023). This behaviour appear to be widespread across animal taxa (Behringer et al., 2018; Lopes et al., 2022). Thus, infection avoidance behaviour has far reaching ecological and evolutionary consequences for hosts and parasites (Buck et al., 2018). Understanding the ecological factors and cues used in driving these behaviours can help better predict the ecology of disease and social interactions (Cable et al., 2017; Esch & Poulin, 1999). This is particularly important given that global changes are expected to modify natural environments at many levels, including the sensory landscape (Dominoni et al., 2020; Rivest et al., 2019), and the transmission of parasitic infections (Krause & Godin, 1996a).

Parasite recognition mechanisms are diverse, but may involve visual cues, such as abnormal host behaviour, visibility of the parasite itself, or physical changes like color patterns, (Behringer et al., 2018; Krause & Godin, 1996; Lopes et al., 2022) and chemical cues, such as pheromones, metabolites and alarm substances (Chivers et al., 2007; Di Bacco & Scott, 2023; Kamio et al., 2022; Stephenson et al., 2018; Yao et al., 2009). In aquatic environments, visual cues are effective for rapid communication, especially in areas where view is unobstructed. However, in complex habitats, such as shallow coastal or littoral zones, visual cues may be limited by physical obstructions (e.g. rugose substrate, branches, aquatic vegetation), water depth, turbidity and/or color (Behringer et al., 2018). Chemical cues, on the other hand, can be transmitted over greater distances in aquatic environments, but can be disrupted or limited by turbulence and anthropogenic inputs such as wastewater and agricultural runoff (Behringer et al., 2018; van der Sluijs et al., 2011). Natural selection should favor organisms that can best exploit the most reliable cues in a given environment (Stephenson et al., 2018). For example, in high turbidity water, chemosensory predators are more abundant and efficient than visual predators at finding prey (Lunt & Smee, 2014, 2015). In dynamic sensory environments, the use of a combination of cues should be favored in order to adjust to temporal variations in the visual and chemical environment (Partan & Marler, 2005; Stephenson et al., 2018). While both sensory mechanisms are useful for parasite recognition and avoidance, their relative importance may vary in different systems or contexts. As anthropogenic activities transform aquatic sensory environments, a better understanding of the cues used to initiate an avoidance response can help predict their impact on fish populations.

The occurrence and the strength of infection avoidance behaviour in hosts can also be influenced by the ecology of the parasite itself, such as its local abundance and its mode of transmission (Moore, 2002). Local parasite abundance can influence learning and the strength of selection for the detection and response to infection risks (Giraldeau & Dubois, 2015; James et al., 2008). In a high parasite load environment, recognition and response to infection cues could thus be enhanced. For instance, fathead minnows (*Pimephales promelas*) naive to *Ornithodiplostomum sp.* parasites do not recognize and avoid visual and chemical free-swimming cercariae cues, whereas experienced fish do (James et al., 2008). In addition, when focusing on the avoidance of conspecifics, the parasite’s mode of transmission may also influence the avoidance response. For parasites that are indirectly transmitted, the selection pressure to avoid infected conspecifics may not be as strong as in directly transmitted infections. Indeed, most studies of infected conspecific avoidance in fishes have focused on host-parasite systems where infection is directly transmitted. However, the presence of infected conspecifics may be a reliable indicator of habitats with a high parasite load and, consequently, a high risk of infection (Karvonen et al., 2004; Krause & Godin, 1996). For example, banded killifish (*Fundulus diaphanus*) prefer uninfected shoals to those infected with the indirectly transmitted trematode *Crassiphiala bulboglossa* causing visible black spot (Krause & Godin, 1996). In addition, conspicuousness of infected individuals resulting from abnormal appearance or behaviour potentially increases group predation risk through the oddity effect (Krause & Godin, 1996; Penry-Williams et al., 2018). Thus avoidance of infected conspecifics may still be advantageous when infection transmission is indirect (e.g. Barber et al., 1998; Krause & Godin, 1996; Tobler & Schlupp, 2007).

Here, we used wild pumpkinseed sunfish (Centrarchidae: *Lepomis gibbosus*, Linnaeus, 1758), a gregarious, common freshwater fish native to eastern North America, to experimentally test for detection and avoidance of infected conspecifics. More specifically, we exposed parasite naive and experienced fish to visual and chemical cues of conspecifics naturally co-infected with indirectly transmitted trematodes (*Uvulifer ambloplitis* and *Apophallus sp.*) and cestodes (*Proteocephalus ambloplitis*). Both of these infections use pumpkinseeds as second intermediate hosts, and are tropically transmitted to final hosts (piscivorous birds or fishes respectively) (Cone & Anderson, 1977; Fischer & Freeman, 1969; Lemly & Esch, 1984). Trematodes encyst in pumpkinseed muscle, under their skin and around fins forming black spots on the body surfaces. These cysts cause skin damage that can release chemical alarm substances (Poulin et al., 1999), and are visually conspicuous, especially at high intensities (Fig. 2.1). Although *P. ambloplitis* cannot be detected visually, infection causes damage to internal organs and is related to altered metabolic rates and cellular processes (Guitard et al., 2022; Mélançon et al., 2023; Thambithurai et al., 2022), which potentially releases detectable chemical cues in their excretions. Both parasites are known to have considerable physiological and behavioural impacts on their hosts (Gradito, 2023; Harrison & Hadley, 1982; Lemly & Esch, 1984; Thelamon, 2023; Tobler & Schlupp, 2007). Sunfish rely on both visual and chemical cues to initiate a range of behavioural responses (Leduc et al., 2003; Marcus & Brown, 2003; Miller, 1963; Stacey & Chiszar, 1978) and have single target acuity (Spratte et al., 2021). Thus, pumpkinseeds may have the ability to perceive and respond to visual and/or chemical cues of infection. At sites where this study took place, there exists a natural variation in infection prevalence and intensity both within and between lakes. Thus, we can test whether the strength of infection avoidance is related to exposure to infected individuals and environments.

Our main objectives were to (1) test the shoaling preferences of uninfected pumpkinseed sunfish based on visual and chemical cues separately. This allowed us to obtain information on the general sociality of pumpkinseeds, which helped us to better address the following aims. Next, we determined: (2) whether uninfected pumpkinseed sunfish display avoidance behaviour towards infected conspecifics, (3) whether pumpkinseeds can detect infection using chemical and/or visual cues, and (4) whether the expression of this behaviour is influenced by the level of parasite exposure of focal fish in their natural environment. To achieve these objectives, we tested parasite naive and parasite experienced populations of pumpkinseeds in visual and chemical binary choice experiments where focal fish had to make a choice in the following three scenarios: (i) uninfected conspecifics vs. lake water with no fish, (ii) infected conspecifics vs. lake water with no fish and (iii) infected vs. uninfected conspecifics. Since pumpkinseeds are gregarious (Holtan, 1998; Vila-Gispert & Moreno-Amich, 2004), tests i and ii served as controls allowing us to establish the degree of sociability of parasite naive and experienced pumpkinseeds populations when given the choice of being with conspecifics or remaining alone. For control tests i and ii, we predicted that (1) pumpkinseeds from both populations would prefer both visual and chemical cues of conspecifics rather than being alone. For test iii, we predicted (2) that pumpkinseeds would actively avoid infected conspecifics and (3) that avoidance would occur in response to both visual and chemical cues, suggesting the presence of infection avoidance behaviour. We also predicted that (4) parasite experienced pumpkinseeds would avoid infected conspecifics more strongly than parasite naive pumpkinseeds. Our results provide insight into the effect of multiple cues on fish infection avoidance response and thus provide a better understanding of the mechanisms underlying this understudied behaviour in fishes.

## MATERIAL AND METHODS

To verify the presence of infected conspecific avoidance behaviour, parasite naive and parasite experienced focal fish were subjected to one of three binary choice treatments: (C1) Control 1 - uninfected conspecifics vs. no fish; (C2) Control 2 - infected conspecifics vs. no fish; or (T) Treatment - infected conspecifics vs. uninfected conspecifics. A single focal fish was exposed to both experiments (visual and chemical), but to a single binary choice treatment, either C1, C2 or T. The order in which the focal fish were subjected to the experiments was randomly selected and focal fish were tested in both experiments (visual and chemical) on different days (1-8 days between tests). For both controls (C1 and C2), 20 naive and 20 experienced focal fish were tested. For the treatment (T), 50 naive and 50 experienced focal fish were tested. C1 and C2 were each completed over the course of four days. The T trials tool 11 days to complete. Experiments were conducted between July 25^th^ and September 15^th^ 2022.

### Fish Collection

Sampling and experiments took place at the Université de Montréal Station de biologie des Laurentides (SBL) in St-Hippolyte, Quebec (45.98898°N – 74.00013°W). Fish were captured in littoral zones of Lake Cromwell (45.59231°N – 73.59565°W) and Lake Triton (45.98769°N – 74,00776°W) between July 9 and August 28, 2022. Both lakes are located in the same drainage basin (Lake Achigan), but are not directly connected, with Lake Triton located upstream of Lake Cromwell. These two populations differ in their prevalence of infection with trematodes and cestodes (Median parasite density: Lake Cromwell = 12.9 trematodes/g, 2.15 cestodes/g; Lake Triton = 0 trematodes/g, 0 cestodes/g) (L.-P. Beauchamp, unpublished data). Lake Cromwell is home to at least eight fish species including the piscivorous smallmouth bass (*Micropterus dolomieu*), which is the final host for cestodes (J. Vignault, unpublished data). Lake Triton has a much simpler fish community with only sunfish and one cyprinid species present and no piscivorous predators (although cannibalism can occur in pumpkinseeds) (J. Vignault, unpublished data).

Focal fish were captured with a seine net. Seine fishing is a non-selective technique that allows sampling of a wide range of behavioural types (Wilson et al., 1993). Stimuli fish used to produce conspecific cues were mostly captured using baited minnow traps and occasionally with seine nets. Minnow traps tend to select for more social and bold individuals (Wilson et al., 1993). Fish were transported to the laboratory in covered, water-filled buckets within 3h of capture.

### Quarantine and Husbandry

Upon arrival at the SBL aquarium facilities, all fish underwent a salt water (3g/L) quarantine treatment for seven days to limit the proliferation of surface infections. The quarantine tanks contained between 50L and 60L of lake water and housed between 20 and 25 fish per tank. Plastic plants and PVC tubes were added to create refuges and reduce aggressive interactions among individuals, and air bubblers ensured oxygen saturation was maintained. Fish were fed once a day (around 8 am) with frozen chironomid larvae (bloodworms). Tanks were cleaned and salt water renewed every day about 3 hours after feeding. After the quarantine treatment, fish were housed in 50.3 × 24.1 × 29, L×W×H cm plastic aquaria (45L of water) in a flow-through system receiving lake water from nearby Lake Croche (45.99206° N – 74.00510° W) also located in the Lake Achigan drainage basin. This water was pumped through a sand filter and an ultraviolet light sterilizer before entering the holding tanks to eliminate any zooplankton. Tanks were cleaned once a day, approximately 3 hours after feeding. The fish experienced approximately 14h light, 10 h darkness, corresponding to an average summer daylight cycle (i.e. 15.7h to 12.2h of daylight; Government of Canada, 2020).

### Focal Fish

Experienced focal fish (n=90) were collected from Lake Cromwell and measured between 5 and 8 cm in standard length (mean SL ± SD: 5.311 ± 0.595 cm, mean mass ± SD: 4.537 ± 1.684 g) and with black spots (Trematoda: *Apophallus* sp. and *Uvulifer* sp.) ranging from 0 - 266 (median: 42). Naïve focal fish (n=90) were collected from Lake Triton and measured between 5 and 8 cm in standard length (mean SL ± SD: 5.598 ± 0.521 cm, mean mass ± SD: 5.217 ± 1.619 g) and with almost no visible black spots (Trematoda: *Apophallus* sp. and *Uvulifer* sp.; min-max: 0 - 1; median: 0). Since males and females are virtually indistinguishable outside of the breeding season (Stacey & Chiszar, 1978), fish sex was determined during post-mortem dissections following experiments (Supplementary tables and figures: Tab. S1). A total of 93 females and 70 males were collected. 17 fish had underdeveloped gonads and could not be sexed. Following salt treatment, all focal fish were subjected to praziquantel treatment (2 mg/L) for 24 h to eliminate cestodes and other unencysted helminths. Following this treatment, fish received a unique color code using visible implant elastomer (VIE tags, Northwest Marine Technology) to identify individuals and their lake of origin. They were also weighed and measured, then transferred to the holding tanks for 6 days before experiments started (14 days following capture). During preliminary trials conducted in September 2021, we found that focal fish fed and responded well to the experimental apparatus as early as 7 days following capture. Focal fish were randomly assigned to different holding tanks (between 10 to 13 fish per tank). Parasite naive and experienced fish were housed in separate tanks and were never in contact throughout the experiments.

**Figure 0.1.**
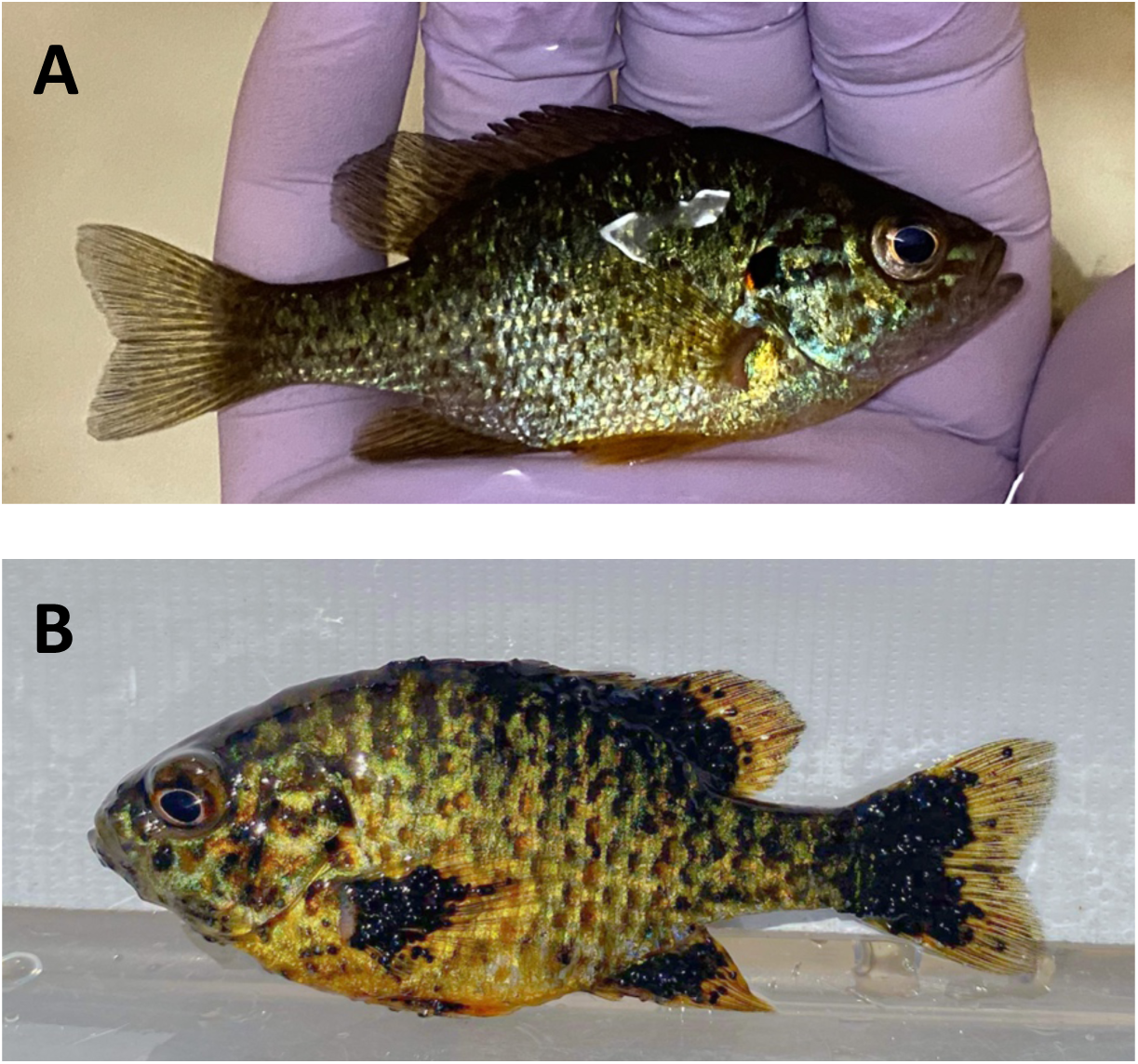
Examples of uninfected and heavily infected stimuli fish. The letters represent the following: (A) Uninfected fish from Lake Triton. (B) Heavily infected fish from Lake Cromwell.

### Stimuli Fish

Infected stimuli fish (n=56: 32 females, 22 males, 2 sex unknown) were collected from Lake Cromwell and measured between 5 and 8 cm in standard length (mean SL ± SD: 5.576 ± 0.605 cm, mean mass ± SD: 5.539 ± 1.943 g) and were heavily infected with black spot (Trematoda: *Apophallus* sp. and *Uvulifer* sp.; min-max: 308 - 1878; median: 727) (Fig. 2.1). These fish were also infected with cestodes, but the exact parasite load was only confirmed during post-mortem dissections following behavioural experiments (Cestoda: *Proteocephalus ambloplites*; min-max: 0 - 114; median: 12). Uninfected stimuli fish (n=56: 28 females, 27 males, 1 sex unknown)) were collected from Lake Triton measuring between 5 and 8 cm in standard length (mean SL ± SD: 5.467 ± 0.297 cm, mean mass ± SD: 4.759 ± 0.840 g) and not infected with black spot (Trematoda: *Apophallus* sp. and *Uvulifer* sp.; min-max: 0 - 0; median: 0) or cestodes (Cestoda: *Proteocephalus ambloplites*; min-max: 0 - 0; median: 0) (Fig. 2.1).

Following salt treatment, stimuli fish were weighed and measured and a visual assessment of blackspot cysts was made to create infected stimuli shoals of seven fish approximately homogeneous in terms of size (mean shoal SL ± SD: 5.521 ± 0.170 cm) and infection intensity (mean ± SD: 767 ± 139 trematodes) (Supplementary tables and figures: Tab. S1). For each binary choice treatment (C1, C2, T), four shoals were created (16 shoals in total. The stimuli shoals were then transferred to the holding tanks for 7 days, for a total habituation period of 14 days before the start of the experiments. Individuals were only ever used in one shoal, and new shoals were used for each treatment (C1, C2 and T) so that the holding time of stimuli fish was similar among treatment and to prevent stimuli shoals from becoming accustomed to the laboratory conditions over time (i.e. 19 days in captivity for both controls and 31 days for the treatment). The stimuli fish did not receive visible elastomer implant or praziquantel treatment. Stimuli fish from the same shoal were housed together in tanks. Within a treatment, stimuli shoals were the same for both visual and chemical experiments. For both C1 and C2, there was one shoal (7 fish) per holding tank. For the Treatment each tank was split in two with a partition to ensure that each stimulus shoal stayed together (14 fish per tank, 7 per section) and accommodate the large number of fish used. Uninfected and infected stimuli shoals were housed in separate tanks and were never in contact throughout the experiments.

### Experiment 1: Visual Cue Choice Tests

#### Experimental apparatus

To measure the behavioural responses of pumpkinseed sunfish to conspecific visual cues, experiments were carried out in a two-compartment choice tank (Fig. 2.2). A glass aquarium (35.5 × 21 × 23 cm, L×W×H) with a water height of 10 cm was used as the choice arena. Two glass aquaria (12 × 21 × 23 cm, L×W×H) placed at opposite ends of the choice arena were used as lateral compartments. These lateral compartments contained either stimulus shoals or lake water with no fish. The water in the experimental devices came from the same source as the holding tanks (i.e. Lake Croche). For C1, an uninfected stimulus shoal was introduced into a lateral compartment and the other compartment contained only lake water.

For C2, an infected stimulus shoal was introduced into a lateral compartment and the other compartment contained only lake water. For T, an infected stimulus shoal was introduced into a lateral compartment and an uninfected shoal stimulus was introduced into the other. There was no water connection among these three aquaria, but focal fish could see the other aquaria through the glass. Removable opaque partitions were inserted between the aquaria. The choice arena, which contained the focal fish, was virtually divided into two choice zones of 11.8 cm in length near the lateral compartments. The choice arena was thus composed of two choice zones near the lateral compartments and a neutral zone in the center. All behavioural trials were video-recorded from above with a digital camera (Go Pro Hero 4) for analysis. Each trial was carried out under constant lighting and temperature (20.4 ± 0.9 °C). To avoid disturbing the fish, the experimental arena was covered with white plastic panels and surrounded by a white curtain. During trials, the experimenter left the room. To avoid visual disturbance, no air bubblers were used during trials.

**Figure 0.2.**
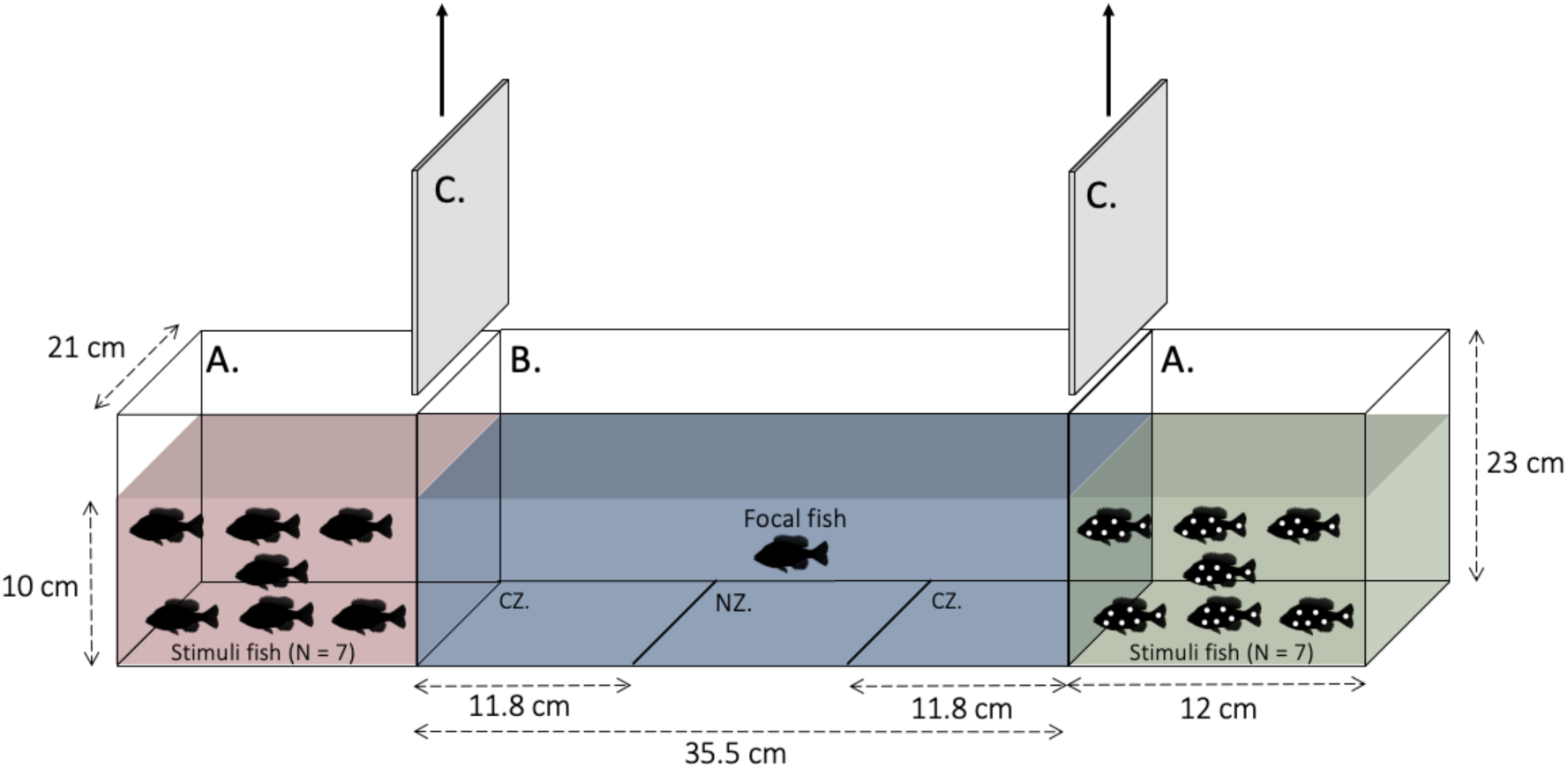
Two-compartment choice tank, front view. There is no water exchange between the three compartments. The letters represent the following: (A) Lateral compartment. (B) Choice arena. (C) Opaque partition. (CZ) Choice zone. (NZ) Neutral zone. Lateral tanks contained infected conspecifics, uninfected conspecifics or no fish, depending on treatment.

#### Protocol

All stimuli fish were fed on the morning of testing. Each day of testing, the same stimulus shoal was used for all trials conducted during the morning and another shoal was used for all afternoon trials. All stimuli shoals were used for the same number of trials within a treatment. The stimulus shoal was placed in the lateral compartment 20 minutes before starting trials. All stimuli fish were transported using a plastic container to limit injury and air exposure and thus limit stress.

All focal fish were fed normally on the morning of testing. Each focal fish was tested individually. Focal fish tested during the same period of time (am or pm) were placed in an 18L bucket (with plants, PVC tubes and bubbler) one hour before the start of the trials to limit stress. Each bucket only contained focal fish from a same holding tank. Fish were transported from the bucket to the experimental device in an opaque glass container filled with the same water as the holding tanks, to minimize stress caused by transport and handling. The focal fish was gently placed in the center of the choice arena without air exposure.

Each test was composed of two trials lasting a total of 20 minutes. The trial began with a 5-minute habituation period during which white opaque plastic partitions prevented visual contact between the focal fish and the lateral compartments for a period of 3 minutes. During the last 2 minutes of habituation, the partitions were removed and the focal fish had access to the lateral compartments with the visual cues to sample the two choice zones. At the end of the habituation period, the trial began and behaviour was recorded. After 5 minutes, the opaque partitions were reinserted, the lateral compartments reversed and the protocol repeated (i.e. 5-minute habituation followed by the second 5-minute trial). The two trials lasted a total of 10 minutes, with the lateral compartments reversed halfway through to avoid lateral bias (Sundin & Jutfelt, 2016). After both trials, the focal fish was put back in its holding tank, the water in the choice arena renewed and an air bubbler placed in each lateral compartment to aerate the water. The order of presentation of the lateral compartments for the first trial was alternated for each fish (e.g. for C1: half the fish had the uninfected stimuli shoal on the right side in trial 1 and the other half on the left side). Between each stimulus shoal, the experimental device was emptied and rinsed. At the end of each binary choice treatment (C1, C2 and T), the entire system was dismantled and washed with water and hydrogen peroxide.

### Experiment 2: Chemical Cue Choice Tests

#### Experimental apparatus

To measure the behavioural responses of pumpkinseed to conspecifics chemical cues, we used a two-current choice flume system (Loligo choice tank system ©) (Fig. 2.3) built with an acrylic reservoir (168 × 40 cm × 25 cm, L×W×H) with a water level of 10 cm. Two parallel laminar currents flowed at a speed of 400 L/h (around 0.27 cm/s) through the tank, which is considerably slower than the average maximum swimming speed of these populations of pumpkinseed sunfish (i.e. 12 to 22 cm/s (J. De Bonville, unpublished data; Brett & Sutherland, 1965)). The flow was constant and precisely adjusted using flowmeters integrated into the system inlet piping. The current speed was chosen to be fast enough to quickly re-establish current separation after a period of active swimming (Atema et al., 2002). The currents, separated by a physical barrier for half of the reservoir, were made laminar using baffles and honeycomb collimator plates installed upstream of the choice arena (Herbert et al., 2011; Jutfelt & Hedgärde, 2013). The choice arena (32 × 40 cm, L×W) contained the focal fish. As suggested by Jutfelt et al. (2017), the compartment size (length and width) was at least 4 times the length of the focal fish (total length from 4.91 cm to 8.80 cm), which is large enough to allow ample movement and reduce confinement stress, but small enough to allow easy access and free movement between the two currents (i.e. both sides of the choice arena). As some focal individuals may demonstrate a side preference independent of the chemical composition of the water (Sundin & Jutfelt, 2016), valves were used to reverse the water currents during the test in order to change the side cues. The valves were located on the inlet piping to avoid disturbance of the focal fish and interruption of the water flow during inversion (Fig. 2.3).

Lateral compartments (51 × 32 × 31 cm, L×W×H) with 37.5 L of water were used to contain the stimulus shoal or lake water with no fish. The water in the experimental devices came from the same source as the holding tanks (i.e. Lake Croche). This water provided standardization, given that neither fish population experienced this water source in the wild (Pérez-Jvostov et al., 2012). For C1, an uninfected stimulus shoal was introduced into a lateral compartment and the other compartment remained with only lake water. For C2, an infected stimulus shoal was introduced into a lateral compartment and the other compartment remained with only lake water. For T, an infected stimulus shoal was introduced into a lateral compartment and an uninfected stimulus shoal was introduced into the other. Both lateral compartments were identical and contained one plastic plant and one PVC tube providing refuges. The water used to create currents was pumped (Eheim Universal 600, Germany) from these compartments to the header tanks of the two-current choice flume. The header tanks were placed 47,0 cm above the reservoir bottom to gravity-fed the inlet piping with water. To ensure constant water volume and pressure to the inlet piping, the header tanks continually overflowed directly in their respective lateral compartment (Jutfelt et al., 2017). During trials, the lateral compartments were constantly supplied with new water, ensuring a continuous inflow of cues at an ecologically relevant concentration into the reservoir. The water was continuously supplied with air through bubblers located both in the head tanks and in the lateral compartments.

All behavioural trials were video-recorded from above with a digital camera (Go Pro Hero 4) for analysis. Each trial was carried out under constant lighting and water temperature (19.5 ± 0.8 °C). To minimize visual disturbance to the focal fish, the reservoir was surrounded by a white curtain. The entire experimental device was also surrounded by a second opaque curtain behind which the experimenter could move.

**Figure 0.3.**
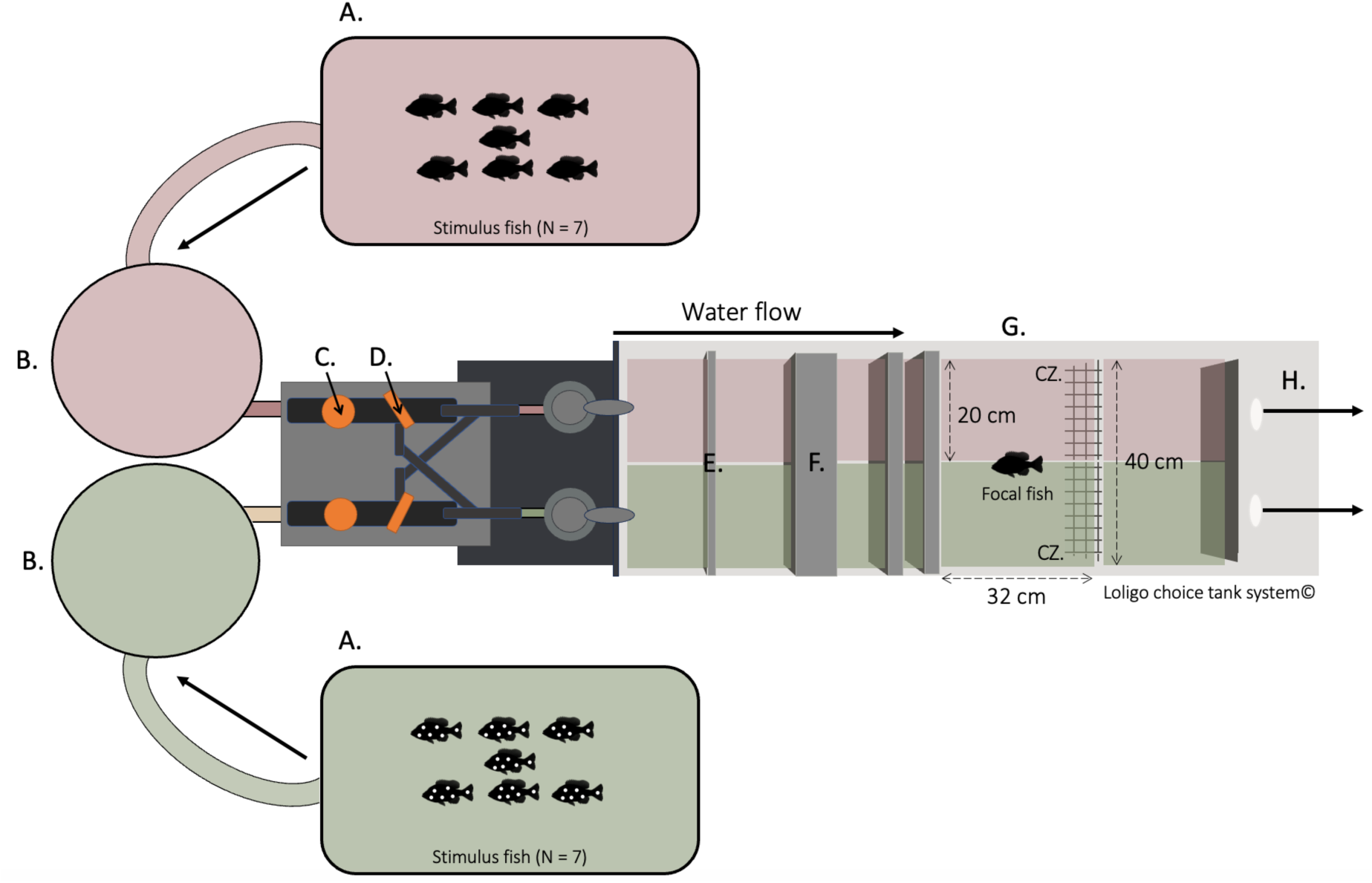
Two-current choice flume, top view. The direction of the arrows indicates the direction of the currents. Letters represent the following: (A) Lateral compartment; (B) Header tank; (C) Flowmeter; (D) Reversing valve; (E) Baffle; (F) Honeycomb collimator plate; (G) Choice arena; (CZ) Choice zone; (H) Acrylic reservoir. Lateral tanks contained infected conspecifics, uninfected conspecifics or no fish, depending on treatment. [Modified from Loligo® Systems (2023)]

#### Protocol

Before each binary choice treatment (C1, C2 and T), a dye test was performed to ensure that currents remained laminar and constant over time (Supplementary tables and figures: Fig. S1). Trials started once the laminar current was reliable (i.e. completely recovered within 30 seconds after a disturbance) (Jutfelt et al., 2017).

The same stimulus shoal was used for all tests conducted on the same day. All stimuli shoals were used for the same number of trials within a treatment. Stimulus fish were fed an hour before moving from their holding tank to the lateral compartments where they were kept overnight (moved between 4 p.m. and 7:30 p.m.) to allow habituation prior to the start of experiments the following morning. All stimuli shoals were transported in a plastic container to limit injury and exposure to air. At the beginning of the test day, the water stream was opened beforehand to renew the water in which the stimulus shoal spent the night, so that the first focal fish did not have a higher concentration of chemical cues.

All focal fish were fed normally on the morning of testing. Focal fish tested during the same period of time (am or pm) were placed in an 18 L bucket (with plants, PVC tubes and air bubbler) one hour before the start of the trials to limit stress caused by capture. Each bucket only contained focal fish from a same holding tank. Focal fish were tested individually. They were moved from the bucket to the choice arena in an opaque glass container filled with the same water as the holding tanks. The focal fish was gently placed in the center of the choice arena without air exposure.

Each test was composed of two sequential trials lasting a total of 20 minutes. The first trial was preceded by a 5-minute habituation period. During the first 3 minutes of the habituation period, the focal fish had no access to chemical cues. During the last 2 minutes, the focal fish had access to the water currents with the chemical cues to enable it to sample both currents. The fish behaviour was recorded over the following 5 minutes. After the first trial, the water currents were reversed and the protocol repeated (i.e. 5-minute habituation followed by the second 5-minute trial). After both trials, the focal fish was put back into its holding tank. Due to logistical constraints, the order of presentation of the lateral compartments for the first trial was the same for all fish (e.g. for C1: all of the fish had the uninfected stimuli shoal on the left side in trial 1). At the end of each day, the stimulus shoal was removed, the system emptied and rinsed. At the end of each binary choice treatment (C1, C2 and T), the entire system was dismantled and washed with water and hydrogen peroxide.

### Euthanasia and Dissection

Once both experiments (visual and chemical) had been completed within a binary choice treatment, all focal and stimuli fish were euthanized with an overdose of eugenol (clove oil). Fish size (total and standard length) and mass were measured immediately after euthanasia. Focal and stimuli fish body condition was estimated using the LeCren’s relative index. This index is used as a body condition factor and is defined as the observed mass of an individual divided by its predicted mass. The latter is determined from the populations’ weight-length relationship. (Gubiani et al., 2020; Le Cren, 1951). The fish were then frozen at -20°C. Frozen fish were transported from SBL to the MIL Campus at the Université de Montréal. Fish dissections were performed between September 28, 2022 and February 2, 2023. During dissections, fish sex was determined. The shoal sex ratio was calculated as the number of males divided by the number of females in each stimulus shoal. Trematodes causing black spot disease were counted only on one half of the fish, i.e. the left side, and the quantity was doubled (De Bonville et al., 2024). The number of trematodes on fish from Lake Triton was not doubled, as only one trematode was exceptionally found in all the fish from this lake. Parasites in fish tissues and body cavity (cestodes, nematodes and yellow grub) were extracted and counted. Parasite density (number/g) was calculated, for both focal and stimuli fish, as the total number of parasites divided by the mass of the fish.

### Videos Analysis

All behavioural trials were analyzed with LoliTrack 5 (Loligo® Systems) automated tracking software. For each trial, video analysis began after the 5-minute habituation period only if the focal fish had visited both choice zones during the last two minutes of this period when the cues were present. For fish that did not visit both zones within the habituation period, behavioural analysis began 10 seconds after the fish had visited both choice zones (Supplementary methods S1). The test time therefore lasted less than 5 min for these fish (from 1min25 to 4min50). Fish that did not visit the two choice zones during the entire test period, for one or both trials, were excluded from the analysis (n= 18). For the visual experiment controls (C1 and C2), 11 fish did not visit both choice zones during the test (i.e. stayed on the side with conspecifics). These fish were included in the analysis as they looked in the direction of the compartment with no conspecifics, and as they visited both sides and showed normal exploratory behavior during the habituation period without visual contact with the lateral compartments. A total of 19 fish for chemical cues and 15 for visual cues were excluded from analysis for various reasons (Supplementary methods S1). The time a focal fish spent in each zone was calculated automatically by the software using the fish’s center of mass. We measured a total of 165 pumpkinseed sunfish for chemical cues (C1: n= 37; C2: n= 38, T: n= 90) and a total of 165 for visual cues (C1: n= 37; C2: n= 37, T: n= 91).

For visual cues, considering that focal fish used the neutral zone as a passage between the two choice zones and did not remain in this zone, the proportion of time spent in the neutral zone (Mean ± SD: 0.079 ± 0.037) was not considered in the analyses.

### Statistical Analysis

All statistical analyses were performed using R 4.2.2 (R Core Team 2022). The proportion of time focal fish spent associating with each cue was calculated as the time spent in each choice zone relative to the time spent in both choice zones. This time proportion served as an attraction index for each cue (Clark et al., 2020; Ehman et Scott, 2001; Rahn et al., 2015). Differences in shoal mean standard length, mass, body condition and parasite density were compared using general linear models two-tailed ANOVAs (Rahn et al., 2018). The normality of residuals was tested using the Shapiro-Wilk normality test and Q-Q plots and the homogeneity of variances was tested using the Bartlett’s test. If either of these assumptions were not met, data were analyzed using nonparametric Kruskal-Wallis tests. There were no significant differences in mean standard length, mass, body condition and parasite density among the stimuli shoals, and therefore these variables were not considered in subsequent analyses (Supplementary tables and figures: Tab. S1).

To test the shoaling preference of focal fish in each experimental condition, we used two-tailed paired Student’s t-tests with the proportion of time spend in each zone as the response variable (Ehman & Scott, 2001; Rahn et al., 2015, 2018). For both controls, we compared the proportion of time spent with conspecific cues vs. no fish cues to assess their preference for shoaling with infected/uninfected conspecifics vs being alone. For the treatment, we compared the proportion of time spent with infected conspecific cue vs uninfected conspecific cue to assess whether focal fish avoid infected conspecifics. Data were tested for normality using the Shapiro-Wilk normality test and Q-Q plots on centered data. Data deviating from normality were arcsine transformed (naive fish in the treatment condition in the chemical cue test) or analyzed using nonparametric paired Wilcoxon signed-ranks test if still deviating from normality after being transformed (naive fish in the treatment condition in the visual cue test).

To examine the impact of the lake of origin on fish attraction to cues, we used generalized linear mixed models (beta error distribution with logit link function in the glmmTMB package; Brooks et al., 2017). For both controls, we used the proportion of time the focal fish spent with conspecific cue (i.e. attraction to conspecific cue) as the response variable. For the treatment, we used the proportion of time the focal fish spent with infected conspecific cue (i.e. attraction to infected conspecific cue) as the response variable. We subtracted a lowercase number (1×10^-12^) to our attraction variable to remove zeros and ones to fit our data into beta distribution limits (Douma & Weedon, 2019). We initially ran full models with the following explanatory variables as fixed effects: the lake of origin, side with cue (Left or Right), trial number (1 or 2), focal fish sex, standard length, body condition, parasite density, holding tank number and time of the experiment (am or pm). We also included the identity, time since eating, sex ratio, mean standard length, mean body condition and mean parasite density of the stimulus shoal. The identity of focal fish (ID) was always included as a random effect. We then used the corrected Akaike information criterion (AICc and ΔAICc) (Hurvich & Tsai, 1989) to compare our models and select the most parsimonious ones that provided significantly better fits to the data. In the end, Our complete models for the visual cues experiment included the lake of origin of the focal fish (to test for differences between naive and experienced fish), the trial number (to control for reversal learning; (Bridgeman & Tattersall, 2019; Lucon-Xiccato & Bisazza, 2014)) and the side of cue (to control for side bias) as fixed effects. Our complete models for the chemical cues experiment included the lake of origin of the focal fish and the trial number only as we could not separate the effect of the trial from the effect of the side. The identity of focal fish (ID) was included as a random effect in all our models. Likelihood ratio tests were performed to assess the significance of the random factor. To do so, we performed an analyse of variance (ANOVA) to compare our complete model to a model without the random factor (ID) (LaHuis & Ferguson, 2009). To calculate confidence intervals of the fixed factors, we used the profile method that computes a profile likelihood with the package MASS (Venables et al., 2002). For our random factor (ID), we calculated confidence intervals of the standard deviation with a Wald t-distribution approximation in the MASS package and then squared it to obtain confidence intervals of the variance parameter.

### Ethical Note

Fish were collected and cared for with approval from the Université de Montréal’s animal care committee and the Ministère de l’Environnement, de la Lutte contre les changements climatiques et de la Faune et des parcs (Comité de Déontologie de l’Expérimentation sur les Animaux; Permit number: 22-025, Collection permit number: 2022-05-16-1971-15-S-P).

## RESULTS

### Experiment 1: Visual Cues for Avoidance of Infected Conspecifics

#### Control 1: Uninfected Conspecific vs. Lake Water

Our statistical analyses included a total of 37 pumpkinseeds (19 naive and 18 experienced). Naive fish spent significantly more time with uninfected conspecifics (M ± SD = 0.682 ± 0.201) than on the side with no fish (M ± SD = 0.294 ± 0.185) (Paired t test: ^*d̅*^ = 0.388, t_37_ = 6.197, *P*< 0.0001) (Fig. 2.4). Similarly, experienced fish spent significantly more time with uninfected conspecifics (M ± SD = 0.710 ± 0.201) than with no fish (M ± SD = 0.268 ± 0.184) (Paired t test: ^*d̅*^ = 0.442, t_35_ = 6.897, *P*< 0.0001) (Fig. 2.4).

We found no significant effect of the lake of origin on time spent with uninfected conspecifics (GLMM: N= 74, b= -0.185 ± 0.366, z= -0.505, df= 6, *P*= 0.613, 95% CI= [-0.922; 0.552]) (Fig. 2.4). The time spent with uninfected conspecifics did vary according to fish ID (GLMM: random factor; N= 74, σ^2^= 0.987, 95% CI= [0.532; 1.780], χ^2^= 55.694, df= 1, *P*_(χ2)_= < 0.001) in our model (Supplementary tables and figures: Tab. S2).

**Figure 0.4.**
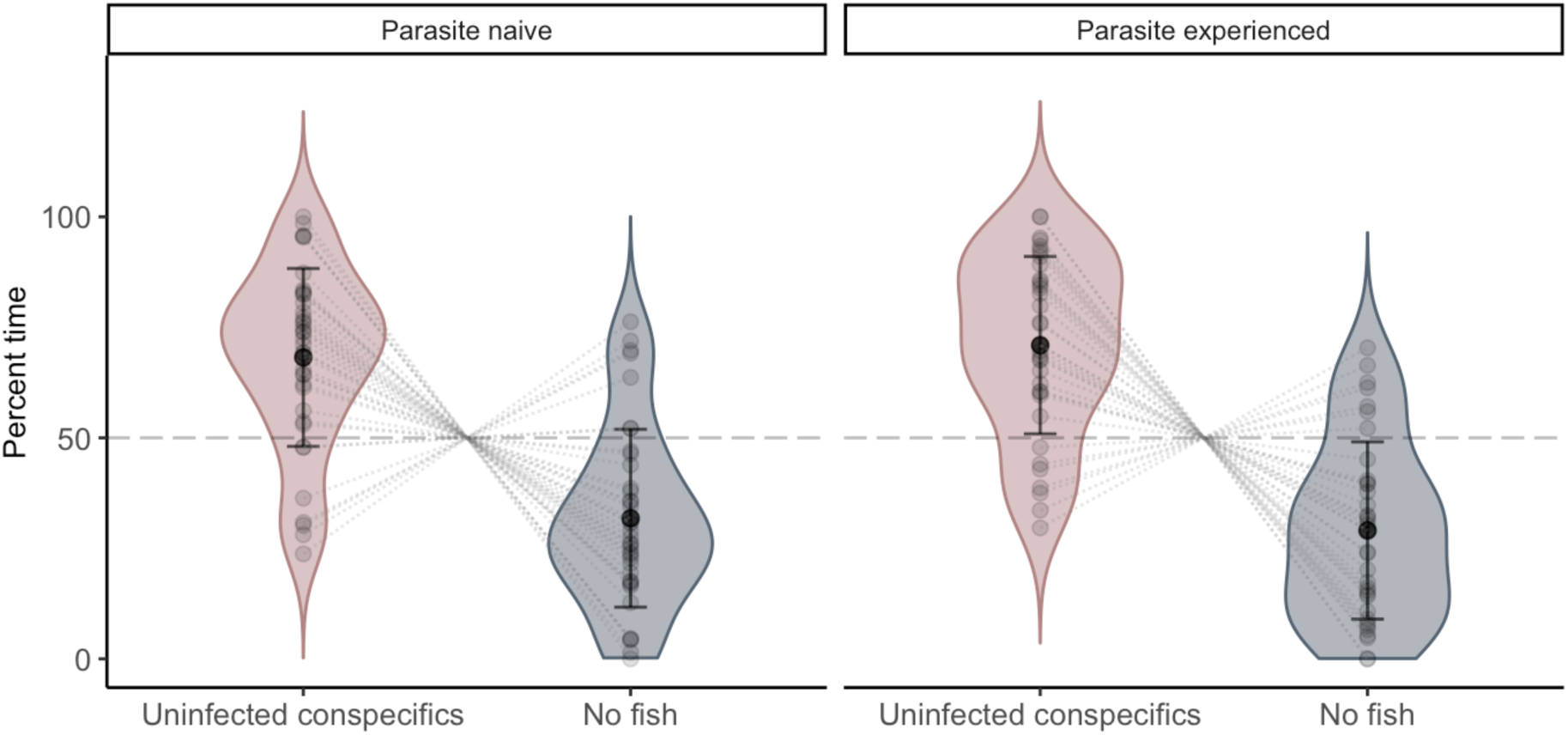
Control 1 – Visual cues: Mean time spent with uninfected conspecifics and no fish visual cues (%) of naive (Trial 1-2, N = 38) and experienced (Trial 1-2, N = 36) pumpkinseed sunfish (Lepomis gibbosus). Dark black dots represent population means; black lines represent standard deviation; pale black dots represent observed data; two pale dots linked by a dotted line represent a single individual. Violon plots are used to show the distribution of the data.

#### Control 2: Infected Conspecifics vs. Lake Water

Our statistical analyses included a total of 37 pumpkinseeds (18 naive and 19 experienced). Naive fish spent significantly more time on the side of the tank with infected conspecifics (M ± SD = 0.773 ± 0.158) than with no fish (M ± SD = 0.227 ± 0.158) (Paired t test: ^*d̅*^ = 0.546, t_35_ = 10.377, *P*< 0.0001) (Fig. 2.5). Similarly, experienced fish spent significantly more time with infected conspecifics (M ± SD= 0.730 ± 0.149) than with no fish (M ± SD = 0.270 ± 0.149) (Paired t test: ^*d̅*^ = 0.459, t_37_ = 9.522, *P*< 0.0001) (Fig. 2.5).

We found no significant effect of the lake of origin on time spent with infected conspecifics (GLMM: N= 74, b= 0.664 ± 0.426, z= 1.558, df= 6, *P*= 0.119, 95% CI= [-0.192; 1.526]) (Fig. 2.5). The time spent with infected conspecifics did vary according to fish ID (GLMM: random factor; N= 74, σ^2^= 1.159, 95% CI= [0.734; 2.456], χ^2^= 64.269, df= 1, *P*_(χ2)_= < 0.001) (Supplementary tables and figures: Tab. S2).

**Figure 0.5.**
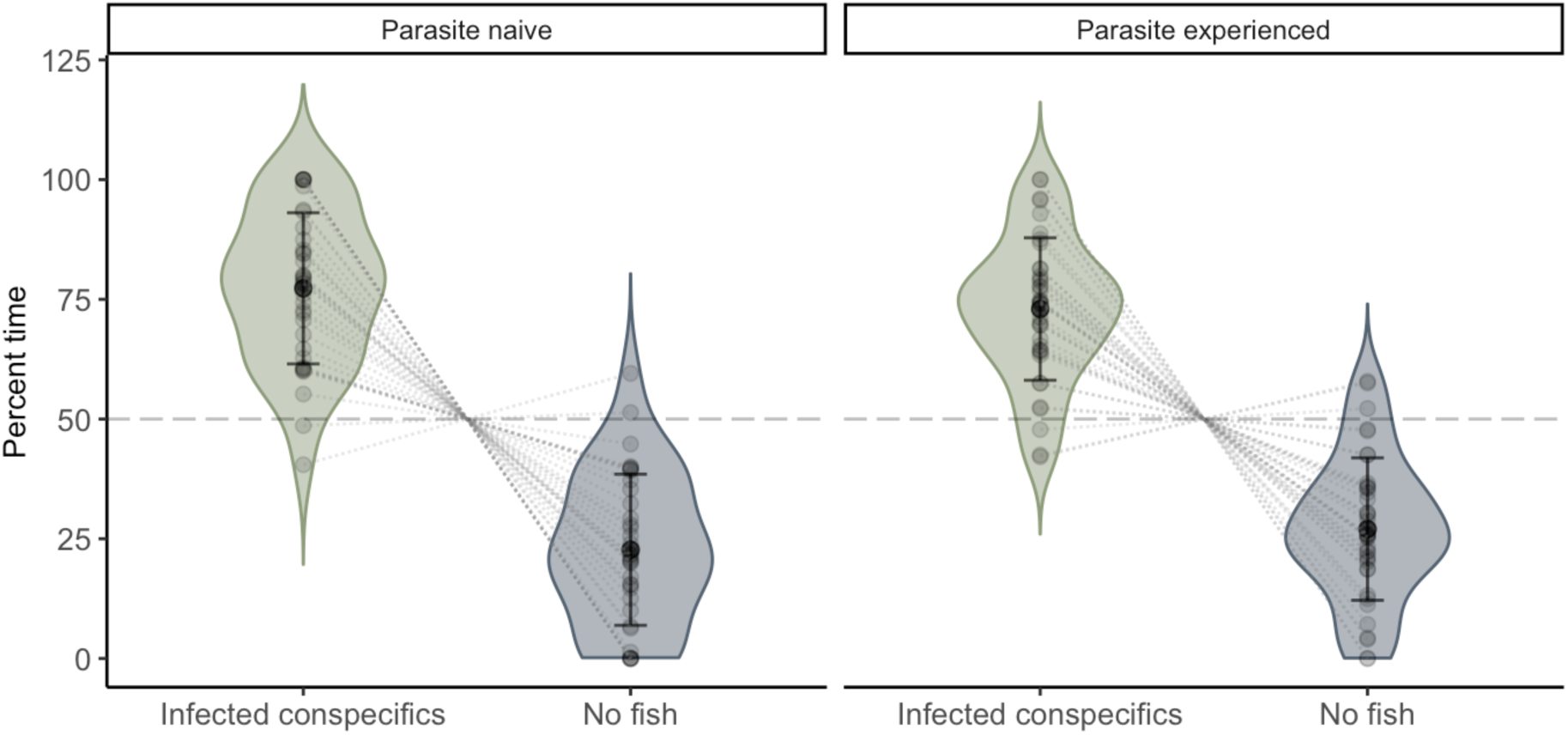
Control 2 – Visual cues: Mean time spent with infected conspecifics and no fish visual cues (%) of naive (Trial 1-2, N = 36) and experienced (Trial 1-2, N = 38) pumpkinseed sunfish (Lepomis gibbosus). Dark black dots represent population means; black lines represent standard deviation; pale black dots represent observed data; two pale dots linked by a dotted line represent a single individual. Violon plots are used to show the distribution of the data.

#### Treatment: Infected vs. Uninfected Conspecifics Cues

Our statistical analyses included a total of 91 pumpkinseeds (47 naive and 44 experienced). For naive fish, data deviated from normality, even after being arcsine transformed, and were thus analyzed using a nonparametric Wilcoxon signed-ranks test. Time spent with uninfected conspecifics (M ± SD = 0.516 ± 0.131) was not different from the time spent with infected conspecifics (M ± SD = 0.484 ± 0.131) (Paired Wilcoxon signed-ranks test: V = 2471, *P*= 0.370) (Fig. 2.6). For experienced fish, time spent with uninfected conspecifics (M ± SD = 0.489 ± 0.148) was not different from the time spent with infected conspecifics (M ± SD = 0.511 ± 0.148) (Paired t test: ^*d̅*^ = -0.022, t_87_ = -0.707, *P*= 0.482) (Fig. 2.6).

We found no significant effect of the lake of origin on time spent with infected conspecifics (GLMM: N= 182, b= 0.099 ± 0.084, z= 1.184, df= 6, *P*= 0.237, 95% CI= [-0.066; 0.265]) (Fig. 2.6). The time spent with infected conspecifics did not vary according to fish ID (GLMM: random factor; N= 182, σ^2^= 2.297e-05, 95% CI= [0; Inf], χ^2^= 0, df= 1, *P*_(χ2)_= 1) (Supplementary Tables: Tab. S2).

**Figure 0.6.**
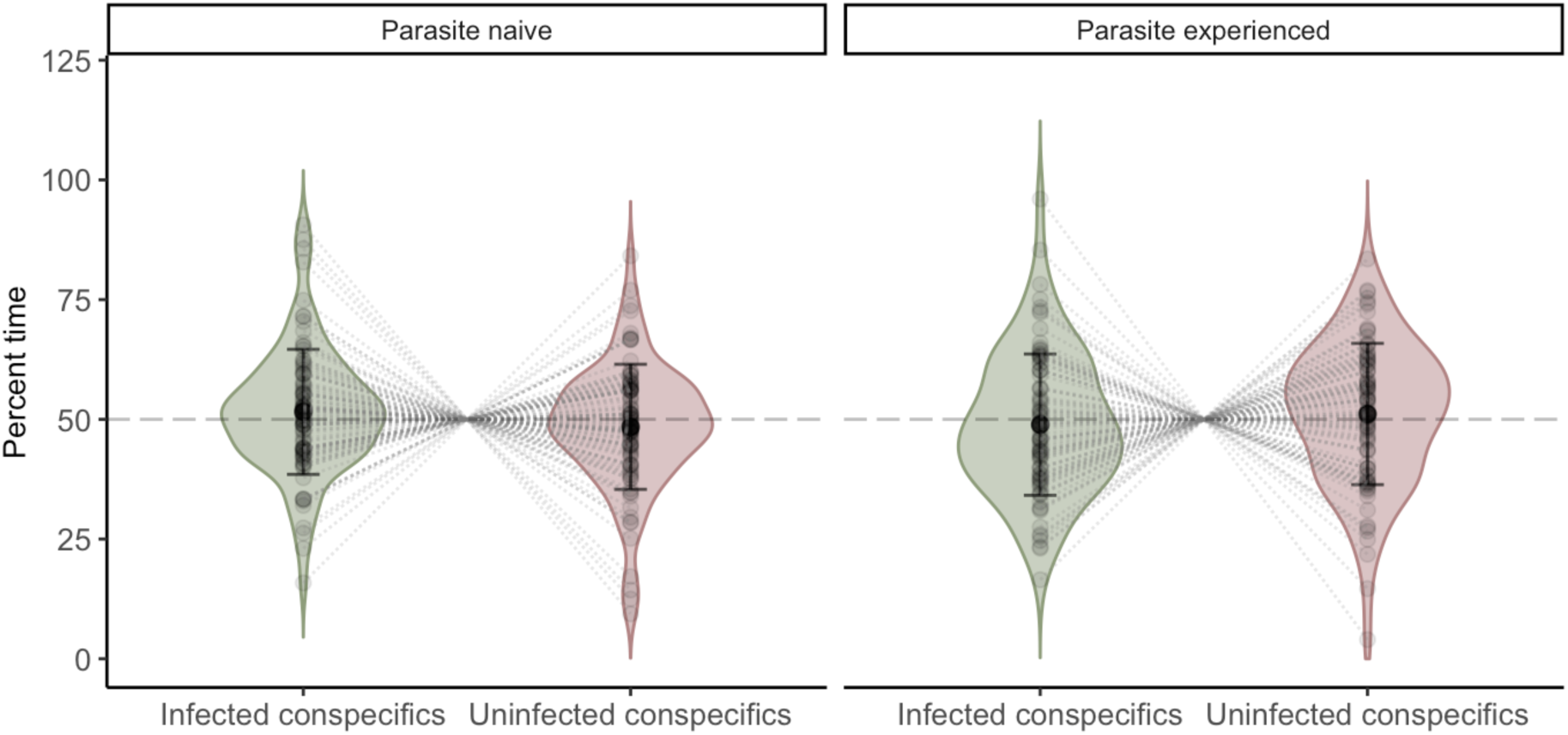
Treatment – Visual cues: Mean time spent with infected conspecifics and uninfected conspecifics visual cues (%) of naive (Trial 1-2, N = 94) and experienced (Trial 1-2, N = 88) pumpkinseed sunfish (Lepomis gibbosus). Dark black dots represent population means; black lines represent standard deviation; pale black dots represent observed data; two pale dots linked by a dotted line represent a single individual. Violon plots are used to show the distribution of the data.

### Experiment 2: Chemical Cues for Avoidance of Infected Conspecifics

#### Control 1: Uninfected Conspecifics Cues vs. Lake Water

Our statistical analyses included a total of 36 pumpkinseeds (18 naive and 18 experienced). Naive fish spent significantly less time with uninfected conspecifics (M ± SD = 0.338 ± 0.199) than with control lake water (M ± SD = 0.662 ± 0.199) (Paired t test: *d̅* = - 0.323, t_35_ = -4.876, *P*< 0.0001) (Fig. 2.7). For experienced fish, the time spent with uninfected conspecifics (M ± SD = 0.441 ± 0.279) was not statistically different from the time spent in control lake water (M ± SD = 0.559 ± 0.279) (Paired t test: *d̅* = -0.117, t_35_ = -1.260, *P*= 0.216) (Fig. 2.7).

We found no significant effect of the lake of origin (GLMM: N= 72, b= -0.427 ± 0.443, z= -0.963, df= 5, *P*= 0.336, 95% CI= [-1.316; 0.463]) (Fig. 2.7). Time spent with uninfected conspecifics did vary according to fish ID (GLMM: random factor; N= 72, σ^2^= 1.084, 95% CI= [0.606; 2.279], χ^2^= 33.6, df= 1, *P*_(χ2)_= < 0.001) (Supplementary tables and figures: Tab. S3).

**Figure 0.7.**
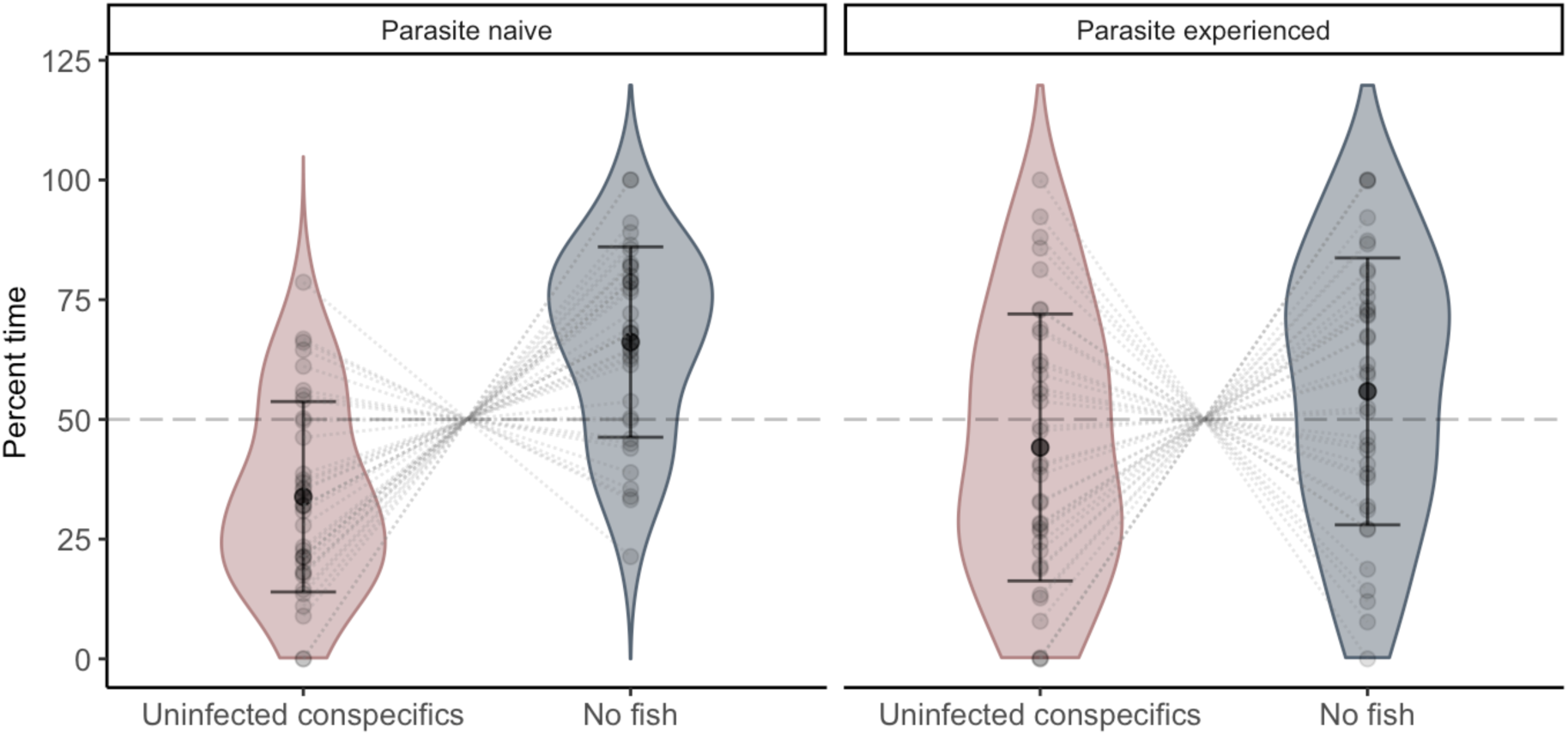
Control 1 – Chemical cues: Mean time spent with uninfected conspecifics and no fish chemical cues (%) of naive (Trial 1-2, N = 36) and experienced (Trial 1-2, N = 36) pumpkinseed sunfish (*Lepomis gibbosus*). Dark black dots represent population mean; black lines represent standard deviation; pale black dots represent observed data; two pale dots linked by a dotted line represent a single individual. Violon plots are used to show the distribution of the data.

#### Control 2: Infected Conspecifics Cues vs. Lake Water

Our statistical analyses included a total of 37 pumpkinseeds (19 naive and 18 experienced). Naive fish spent significantly less time with infected conspecifics (M ± SD = 0.432 ± 0.165) than with control lake water (M ± SD = 0.568 ± 0.165) (Paired t test: *d̅* = - 0.135, t_37_ = -2.533, *P*= 0.016) (Fig. 2.8). Experienced fish also spent significantly less time with infected conspecifics (M ± SD = 0.356 ± 0.136) than with control lake water (M ± SD = 0.644 ± 0.136) (Paired t test: *d̅* = -0.288, t_35_= -6.363, *P*< 0.0001) (Fig. 2.8).

We found a significant effect of the lake of origin on time spent with infected conspecifics (GLMM: N= 74, b= 0.300± 0.146, z= 2.053, df= 5, *P*= 0.040, 95% CI= [0.010; 0.589]). Naive fish spent significantly more time with infected conspecifics than experienced fish (Fig. 2.8). Time spent with infected conspecifics did not vary according to fish ID (GLMM: random factor; N= 74, σ^2^= 2.155e-05, 95% CI= [0; Inf], χ^2^= 0, df= 1, *P*_(χ2)_= 1) (Supplementary tables and figures: Tab. S3).

**Figure 0.8.**
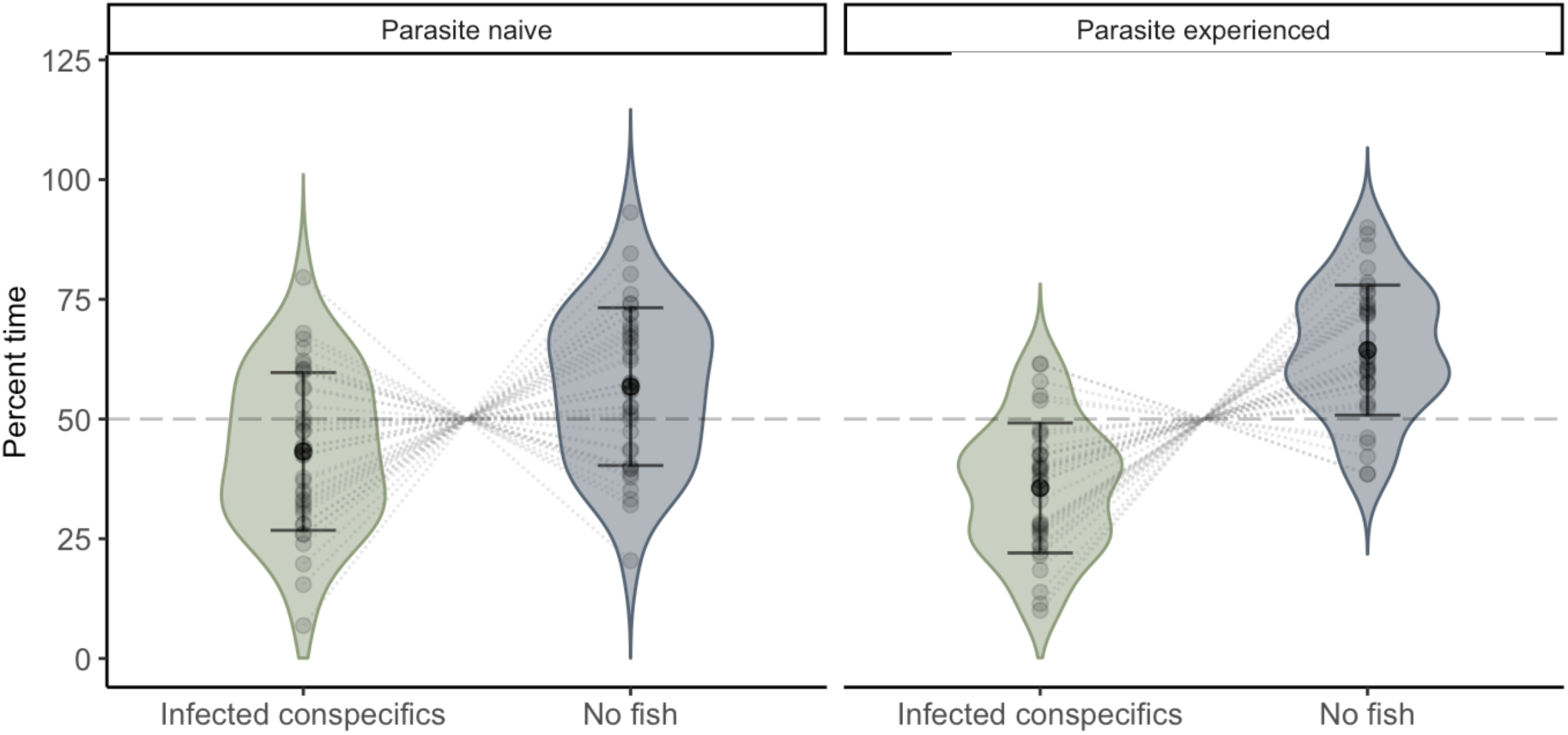
Control 2 – Chemical cues: Mean time spent with infected conspecifics and no fish chemical cues (%) of experienced (Trial 1-2, N = 38) and naive (Trial 1-2, N = 36) pumpkinseed sunfish (Lepomis gibbosus). Dark black dots represent population means; black lines represent standard deviation; pale black dots represent observed data; two pale dots linked by a dotted line represent a single individual. Violon plots are used to show the distribution of the data.

#### Treatment: Infected vs. Uninfected Conspecifics Cues

Our statistical analyses included a total of 88 pumpkinseeds (41 naive and 47 experienced). For naive fish, data deviated from normality and were arcsine transformed. Naive fish spent significantly less time with infected conspecific (M ± SD = 0.577 ± 0.272) than with uninfected conspecific cues (M ± SD = 0.423 ± 0.272) (Paired t test on arcsine transformed data: *d̅* = -0.180, t_81_ = -2.498, *P*= 0.015) (Fig. 2.9). For experienced fish, there was no difference in the time spent with infected conspecific (M ± SD = 0.469 ± 0.235) compared to uninfected conspecific cues (M ± SD = 0.531 ± 0.235) (Paired t test: *d̅* = -0.062, t_93_ = -1.286, *P*= 0.202) (Fig. 2.9).

We found no significant effect of the lake of origin on time spent with infected conspecifics (GLMM: N= 175, b= -0.166 ± 0.234, z= -0.711, df= 5, *P*= 0.477, 95% CI= [- 0.631; 0.295]). (Fig. 2.9). The time spent with infected conspecifics did vary according to fish ID (GLMM: random factor; N= 176, χρ^2^= 0.846, 95% CI= [0.443; 1.158], χ^2^= 56.283, df= 1, *P*_(χ2)_ < 0.0001) (Supplementary tables and figures: Tab. S3).

**Figure 0.9.**
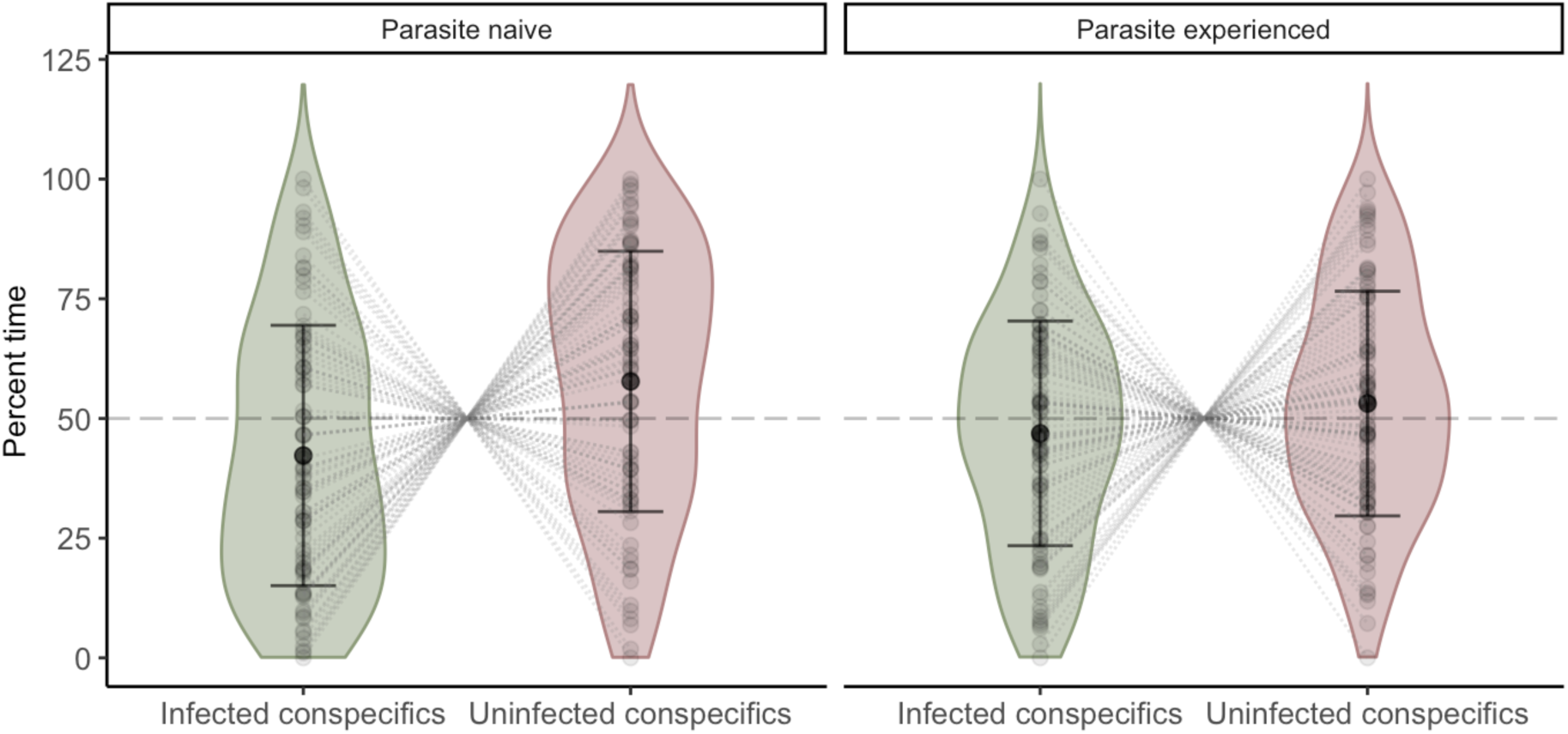
Treatment – Chemical cues: Mean time spent with infected conspecifics and uninfected conspecifics chemical cues (%) of naive (Trial 1-2, N = 94) and experienced (Trial 1-2, N = 82) pumpkinseed sunfish (Lepomis gibbosus). Dark black dots represent population means; black lines represent standard deviation; pale black dots represent observed data; two pale dots linked by a dotted line represent a single individual. Violon plots are used to show the distribution of the data.

## DISCUSSION

We experimentally tested for visual and chemical detection and avoidance of infected conspecifics on parasite naive and experienced pumpkinseeds. We found that pumpkinseeds preferred to shoal with conspecifics over being alone when given visual cues, but avoided conspecifics and preferred to remain alone when given chemical cues regardless of the infection status of their conspecifics. This suggests that, in natural environments, chemical and visual cues are not redundant, and that pumpkinseeds rely on both to make relevant social decisions. We also found significant inter-individual variation in terms of shoaling preference in the presence of chemical cues whereas this variation was not observed in the presence of visual cues. Additionally, we show that the parasite-naive population avoids the chemical cues of infected conspecifics whereas the parasite-experienced population does not. Together, these findings suggest that pumpkinseeds use chemical rather than visual cues to discriminate between infected and uninfected conspecifics and that they can better evaluate infection risk with chemical cues than with visual cues. We also suggest that parasite experienced pumpkinseeds have become habituated to infection cues of conspecifics compared to parasite naive pumpkinseeds that expressed an avoidance response. Our results highlight the importance of considering different sensory cues as well as indirectly transmitted parasites when studying infection avoidance behaviour and shoaling decisions in fishes.

### Sunfish use visual cues for social behaviour

Pumpkinseed sunfish are a gregarious species that use sight to engage in behaviours such as mating, predation and agonistic interactions (Blanckenhorn, 1992; Kieffer & Colgan, 1991; Miller, 1963; Spratte et al., 2021; Stacey & Chiszar, 1978; Vila-Gispert & Moreno-Amich, 2004). Based on these characteristics, we first established individual social preference in a binary choice experiment as a proof of concept to evaluate pumpkinseeds’ use of visual cues in choosing shoal mates. As predicted, in both control tests (C1 and C2), pumpkinseeds spent more time with conspecifics than alone, and this was regardless of conspecific infection status or whether the focal fish was from parasite experienced or naive populations. (Fig. 2.4 and 2.5). These results suggest that focal fish can recognize both populations as conspecifics and strongly respond to the visual cues by making a clear choice. However, trial number had a slight negative effect on time spent with conspecifics with less strong preference for shoaling with conspecifics on the second trial (Supplementary tables and figures: Tab. S2). Nevertheless, these results do not affect our conclusion as focal fish showed a clear preference for conspecifics visual cues in the second trial of both controls (Supplementary Results). These results confirmed that our experimental design was appropriate for assessing avoidance behaviour in this species.

When looking at focal fish preference between infected and uninfected conspecific visual cues (T), we found that the average proportion of time spent associating with both types of conspecifics was not different (Fig. 2.5). This response did not differ between individuals. This result held regardless of whether focal fish were from parasite naive or experienced populations. Naive pumpkinseeds were collected from lakes that are not infected with these species of parasites, and thus would not have learned to recognize nor respond to the cues (James et al., 2008). Considering that experienced fish showed the same response as naive ones, we suggest that pumpkinseeds do not use visual cues to discriminate between shoal mates infected with black spot disease. Previous work has shown that pumpkinseeds use a variety of visual cues and can identify conspecifics and related species up to a few meters away (Kieffer & Colgan, 1991; Miller, 1963; Spratte et al., 2021; Stacey & Chiszar, 1978). Consequently, they likely have the ability to see blackspots (i.e. encysted trematodes) as well as other morphological or behavioural differences between infected and uninfected individuals. We therefore suggest that, even if pumpkinseeds can perceive visual infection cues, it may not elicit an avoidance response because it is not a reliable indicator of risk.

Parasites in our study system are not directly transmitted by contact between individuals (Cone & Anderson, 1977; Fischer & Freeman, 1969; Lemly & Esch, 1984) and the selection pressure to detect and avoid conspecific infection cues may not be as strong as in directly transmitted infections. Nevertheless, our results contrast with similar two-choice experiments where focal fish avoided conspecifics infected with indirectly transmitted trematodes (e.g. Krause & Godin, 1996; Tobler & Schlupp, 2007). Considering that the infected pumpkinseeds from our study system are host to several other non-visible endoparasites (De Bonville et al., 2024; Guitard et al., 2022; Mélançon et al., 2023; Thambithurai et al., 2022), visual cues of trematode infection may not be a reliable indicator of a conspecific health, nor of high-risk habitats. For instance, when cannibalism occurs, pumpkinseeds can serve as final hosts to adult cestodes, i.e bass tapeworms in our study system (V. Mélançon unpublished data; Lai & Johnston, 2002). This is not possible for trematodes, as their final hosts are piscivorous birds (Roberts & Janovy, 2005). Given that avoidance behaviours are risk-sensitive (Buck et al., 2018; Stephenson et al., 2018), visual cues may not be a sufficiently reliable indicator of risk to depend on in terms of shoaling preference and infection avoidance when fish are co-infected with parasites requiring different final host to complete their lifecycle.

### Sunfish use chemical cues for risk assessment

Contrary to our prediction, focal fish generally spent more time alone than with either infected or uninfected conspecifics when given chemical cues (Fig. 2.6 and 2.7). The response varied slightly between the two populations. Compared to naive focal fish, experienced focal fish avoided uninfected conspecifics in C1 less strongly and infected conspecifics in C2 more strongly. However, these differences were small (p-values > 0.04, estimates < 0.5). Overall, this suggests that both parasite naive and experienced focal fish were able to receive and respond to chemical cues in our experimental set-up, which is consistent with studies on the use of chemical cues in pumpkinseeds (Leduc et al., 2003; Marcus & Brown, 2003; Xia et al., 2018). Compared to the results obtained for visual cues (C1 and C2) and studies on pumpkinseed sociality (Blanckenhorn, 1992; Vila-Gispert & Moreno-Amich, 2004), these avoidance responses are opposite to our predictions that both populations would prefer chemical cues of conspecifics rather than being alone. We suggest that visual and chemical cues are not redundant (i.e. do not elicit the same response when presented separately; (Partan & Marler, 2005)) for pumpkinseeds, and that they need both to assess the risks and make a relevant social decision.

When looking at the preference between infected and uninfected conspecific chemical cues (T), we found that naive fish spent, on average, less time with infected conspecific cues, whereas experienced fish spent similar average times with both cues (Fig. 2.8). This suggests that parasite naive fish have a greater tendency to avoid the chemical cues of infected conspecifics than experienced fish. However, the difference between the two populations was not significant, which limits the conclusions that can be drawn, but opens the door to further research into the role of learning in the recognition of infection cues, which remains poorly understood (Klemme & Karvonen, 2016; Wisenden et al., 2009). Indeed, our results contrast with studies in which parasite experienced individuals responded more strongly to infection cues (James et al., 2008; Keymer et al., 1983; Klemme & Karvonen, 2016). In our study, the lack of a strong avoidance of infected conspecifics in experienced fish could be a case of habituation (Giraldeau & Dubois, 2015). When encysting, trematodes cause skin damage that may release alarm substances (Cone & Anderson, 1977; Happel, 2019; Poulin et al., 1999) and other chemical cues. Cestodes cause damage to internal organs and affect metabolic processes (Guitard et al., 2022; Mélançon et al., 2023; Thambithurai et al., 2022), which also potentially releases detectable chemical cues in their excretions. These chemical cues can be interpreted as an indication of danger in the parasite naïve population. Considering that 90.0% of the parasite experienced population was infected with trematodes and 96.2% with cestodes (Beauchamp, in prep; Vigneault, in prep), experienced pumpkinseeds may be habituated to these cues. Thus they may not strongly avoid infection cues since avoiding infected conspecifics and habitat in the wild may be difficult to do when infection prevalence is so high. However, our inferences are limited by the fact we do not know the exact chemical cues involved. Further studies on the exact composition of these chemical cues are needed.

While considering the significantly different avoidance response between the two populations, there was also a large variability in the individual responses of focal fish to conspecific chemical cues with some individuals from both populations strongly avoiding, strongly preferring, or showing no preference (Fig. 2.9). This response differs from the visual cues experiment treatment (T), where there was no distinction among individuals, averaging around 50%, thus indicating no specific choice (Fig. 2.6). We suggest that individuals make different decisions in terms of shoaling preferences based on chemical information. Shoaling preferences can be influenced by factors such as the ability to compete with conspecifics for resources and protection against predators (Krause & Godin, 1994, 1996; Metcalfe & Thomson, 1995; Rahn et al., 2018). These factors can be influenced by the intensity and stage of infection of conspecifics, reducing their competitiveness for foraging and increasing their predation risk (Krause & Godin, 1994, 1996; Metcalfe & Thomson, 1995; Rahn et al., 2018). Given that chemical cues provide reliable public information (James et al., 2008; Marcus & Brown, 2003), it may be possible for pumpkinseeds to evaluate the level of risk and the optimal shoaling strategy for them. These two strategies (avoid or prefer infected shoals) support the presence of trade-offs for pumpkinseeds between foraging, predation and potentially infection risk when choosing shoalmates. This is consistent with a study which found evidence for behavioural syndromes between activity and exploration in uninfected pumpkinseeds (Gradito, 2023) suggesting a trade-off between foraging and predation. While keeping in mind that the avoidance response differs between the two populations, the possible capacity of detecting and evaluating the health status of conspecifics seems to be innate as it held regardless of whether fish were from parasite naive or experienced populations.

## Conclusion

Many studies show that pumpkinseeds use visual cues for social behaviours such as recognition, mating and agonistic interactions (Kieffer & Colgan, 1991; Miller, 1963; Spratte et al., 2021; Stacey & Chiszar, 1978). Studies carried out in the context of chemical cues tend to focus on predator avoidance and local risk assessment using conspecific and heterospecific alarm substances (Leduc et al., 2003; Marcus & Brown, 2003; Xia et al., 2018). Here, by measuring the response to different communication cues separately, we provide evidence that pumpkinseeds use both visual and chemical cues to make relevant shoaling decisions and that both cues are not redundant, providing increased information content per unit of time for this species (Partan & Marler, 2005). Visual cues seem to be used for conspecific recognition whereas chemical cues seem to be for risk assessment. Considering that both types of communication are essential for social behaviours, pumpkinseeds may be impacted by the modification of sensory landscapes due to global changes. Indeed, the detection of chemical cues in pumpkinseeds can be altered by water acidity (Leduc et al., 2003) and their visual acuity is influenced by the physical characteristics of the environment (Spratte et al., 2021). By altering the chemical and visual landscapes of waterbodies, the threats posed by acid rain, browning and eutrophication of freshwater ecosystems is a growing concern to researchers interested in sensory ecology, and the implications of such changes to disease ecology warrants further investigation (Dominoni et al., 2020; Mucci et al., 2011; Rivest et al., 2019; Schindler, 1988; Vander Zanden & Vadeboncoeur, 2020).

## ACKNOWLEDGEMENT

We acknowledge the traditional lands of the Kanien’kehá:ka, Omàmiwinini, and Anishinabewaki First Nations on which the field and laboratory work for this project took place. We thank Gabriel Lanthier, Georges-Étienne Charette and Gilles Côté for their help with logistic and assembly of experimental set-up. We also thank Jeffrey D’amour Pigeon, Andréa Serres, Maryane Gradito, Juliane Vigneault and Laurie Provençal for assistance on field and during experiments; Aya Maria Bouyarden, Félix Lamarche and Matthew Archambault for help with dissections and video analysis; Renata Mazzei, Jérémy De Bonville, Marie Levet and Vincent Mélançon for helpful comments on the study design and on data interpretation.

## APPENDIX

### Supplementary Methods

#### S1. Modification and exclusion from analyses

##### Experiment 1: Visual cue choice tests

The number of fish that visited the two choice zones after the habituation period were the following: two focal fish for C1 (two naive); six focal fish for C2 (three naive and three experienced); and 14 focal fish for T (eight naive and six experienced). The test time therefore lasted less than 5 minutes for these fish. 15 trials were excluded from analysis. For C1, three focal fish (one naive and two experienced) were excluded from analysis because they were sick (i.e. vomited and ate it). For C2, three focal fish (two naive and one experienced) were excluded from analysis: two because they were sick (i.e. vomited and ate it) and one because the video was not analyzable. For T, nine focal fish (three naive and six experienced) were excluded from analysis: three because they did not visit both choice zones during one or both trials, five because there were problems during the video analysis (i.e. software did not detect the fish), one because a video was lost due to equipment failure. For the visual experiment controls (C1 and C2), a total of 11 fish did not visit both choice zones during the test (i.e. stayed on the side with conspecifics) but were analyzed. For C1, a total of six fish (two naive and four experienced). For C2, a total of five fish (three naive and two experienced).

##### Experiment 2: Chemical cue choice tests

The number of fish that visited the two choice zones after the habituation period were the following: three focal fish for C1 (one naive and two experienced); six focal fish for C2 (four naive and two experienced); and 11 focal fish for T (six naive and five experienced). The trial time therefore lasted less than 5 minutes for these fish. For C1, four focal fish (two naive and two experienced) were excluded from analysis: three because they did not visit both choice zones during one or both trials, and one because there were problems during the trial (i.e. the focal fish was put back in its holding tank between the two trials). For C2, three focal fish (one naive and two experienced) were excluded from analysis because they did not visit both choice zones during one or both trials. For T, 12 focal fish (nine naive and three experienced) were excluded from analysis: nine because they did not visit both choice zones during one or both trials, and one because there were problems during the trial (i.e. the camera stopped working and the focal fish saw the experimenter).

### Supplementary Tables and Figures

**Figure S1.**
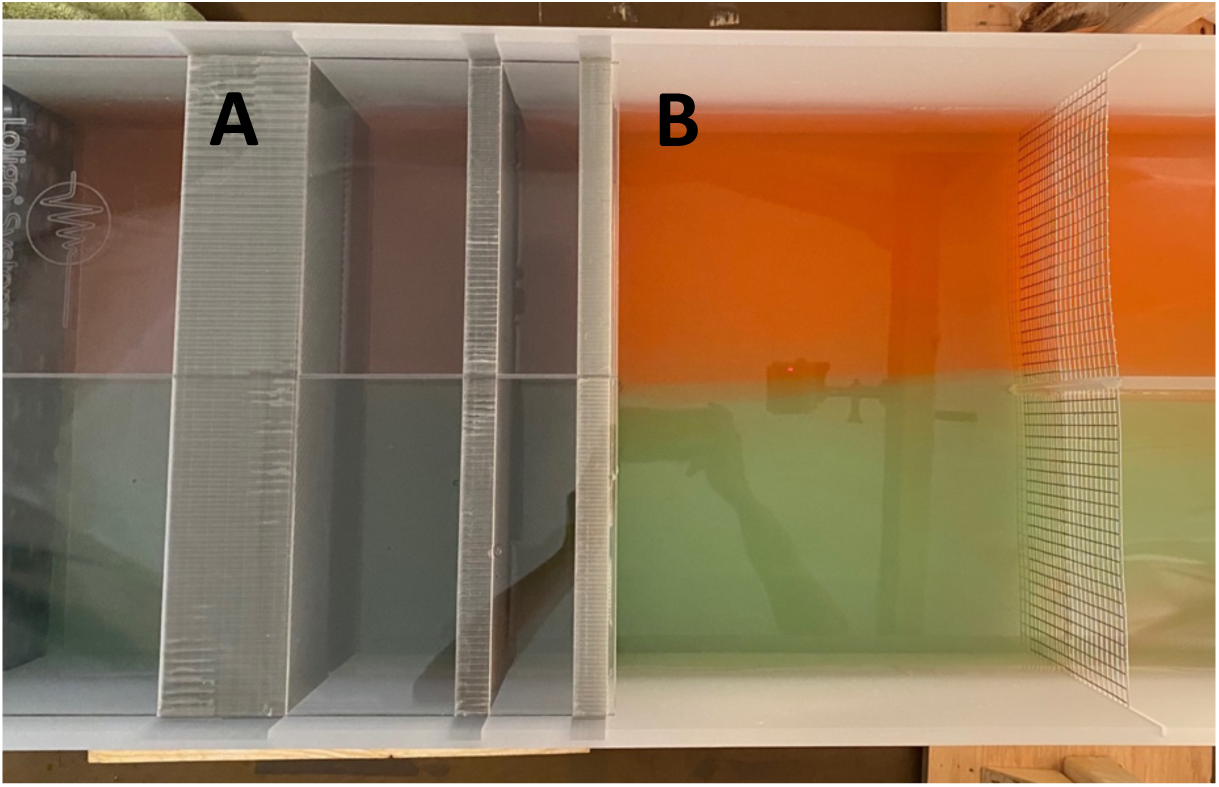
Dye test of the two-current choice flume, top view. Letters represent the following: (A) Honeycomb collimator plate; (B) Choice arena. The two currents, represented by green and red dye, do not mix in the choice arena.

**Table S1.** Characteristics of the stimuli shoals used for each binary choice treatment. For each treatment (C1, C2 and T), new individuals were used to create the stimuli shoals of conspecifics. Each of them consisted of seven stimuli fish (n=7 per shoal). For each shoal, abbreviations represent the following: (Mass) Mean mass ± Standard deviation. (SL) Mean standard length ± Standard deviation. (Trematodes) Median; min-max. (Cestodes) Median; min-max. For information on both parasites, the median is used to compensate for the size and mass of each fish. (Sex) Number of females; Number of males; Number of unknown sex.

| Shoal number | Mass | SL | Trematodes | Cestodes | Sex |
| --- | --- | --- | --- | --- | --- |
| **C1 - Uninfected conspecifics (n = 28)** |  |  |  |  |  |
| 1 | 4.94±0.38 | 5.55±0.11 | 0; 0-0 | 0; 0-0 | 5; 1; 1 |
| 2 | 4.49±0.51 | 5.39±0.20 | 0; 0-0 | 0; 0-0 | 5; 2; 0 |
| 3 | 4.89±1.29 | 5.46±0.42 | 0; 0-0 | 0; 0-0 | 3; 4; 0 |
| 4 | 4.46±0.77 | 5.30±0.26 | 0; 0-0 | 0; 0-0 | 4; 3; 0 |
| Comparison | Kruskal-Wallis<br>$H_3=2.33, P=0.51$ | Kruskal-Wallis<br>$H_3=3.67, P=0.30$ | NA | NA | NA |
| **C2- Infected conspecifics (n = 28)** |  |  |  |  |  |
| 1 | 5.47±1.73 | 5.59±0.51 | 1004; 510-1878 | 9; 2-33 | 4; 2; 1 |
| 2 | 6.07±2.28 | 5.79±0.61 | 552; 308-1116 | 8; 0-30 | 6; 1; 0 |
| 3 | 6.41±2.65 | 5.77±0.78 | 746; 390-1218 | 15; 3-37 | 4; 3; 0 |
| 4 | 6.21±1.99 | 5.90±0.66 | 600; 504-1566 | 10; 2-43 | 2; 4; 1 |
| Comparison | ANOVA | ANOVA | Kruskal-Wallis | Kruskal-Wallis | NA |
| | $F_{1,26}=0.5, P=0.49$ | $F_{1,26}=0.74, P=0.40$ | $H_3=4.96, P=0.17$ | $p=0.862$ | |
| **T - Uninfected conspecifics (n = 28)** |  |  |  |  |  |
| 1 | 4.75±1.05 | 5.49±0.39 | 0; 0; 0-0 | 0; 0; 0-0 | 3; 4; 0 |
| 2 | 4.46±0.64 | 5.31±0.21 | 0; 0; 0-0 | 0; 0; 0-0 | 4; 3; 0 |
| 3 | 5.07±0.99 | 5.69±0.32 | 0; 0; 0-0 | 0; 0; 0-0 | 2; 5; 0 |
| 4 | 5.01±0.87 | 5.54±0.27 | 0; 0; 0-0 | 0; 0; 0-0 | 2; 5; 0 |
| Comparison | **ANOVA** | **ANOVA** | **NA** | **NA** | **NA** |
| | $F_{1,26}=0.85, P=0.36$ | $F_{1,26}=0.97, P=0.33$ | | | |
| **T - Infected conspecifics (n = 28)** |  |  |  |  |  |
| 1 | 5.10±2.33 | 5.36±0.70 | 602; 508-964 | 13; 5-25 | 6; 1; 0 |
| 2 | 5.30±1.53 | 5.42±0.54 | 748; 448-1506 | 11; 0-20 | 5; 2; 0 |
| 3 | 4.83±1.75 | 5.30±0.62 | 756; 620-902 | 13; 7-114 | 2; 5; 0 |
| 4 | 4.90±1.22 | 5.46±0.31 | 642; 494-526 | 12; 9-19 | 3; 4; 0 |
| Comparison | **Kruskal-Wallis** | **ANOVA** | **Kruskal-Wallis** | **Kruskal-Wallis** | **NA** |
| | $H_3=0.80, P=0.85$ | $F_{1,26}=0.03, P=0.86$ | $H_3=4.12, P=0.25$ | $H_3=2.07, P=0.56$ | |

**Table S2.**
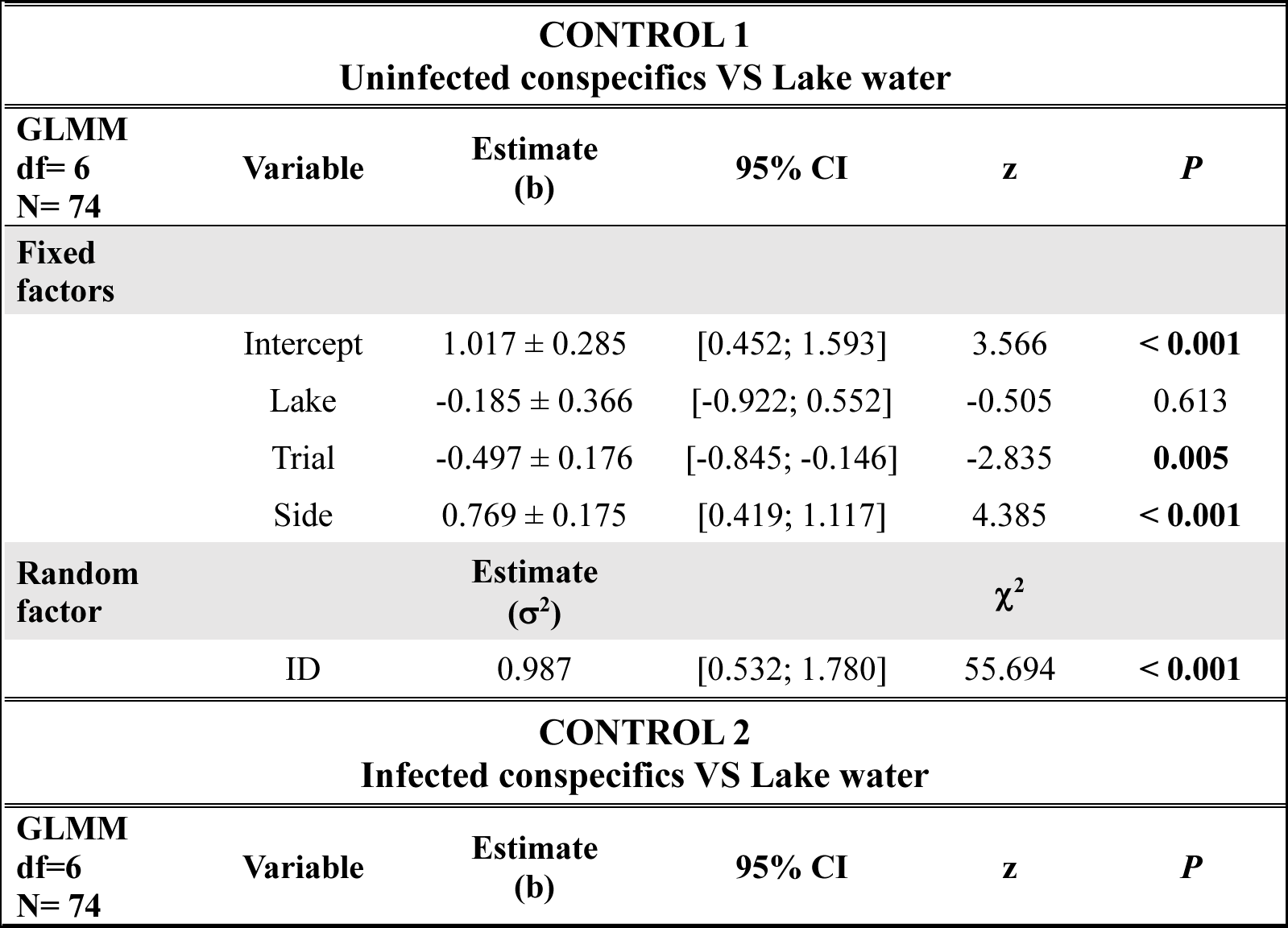
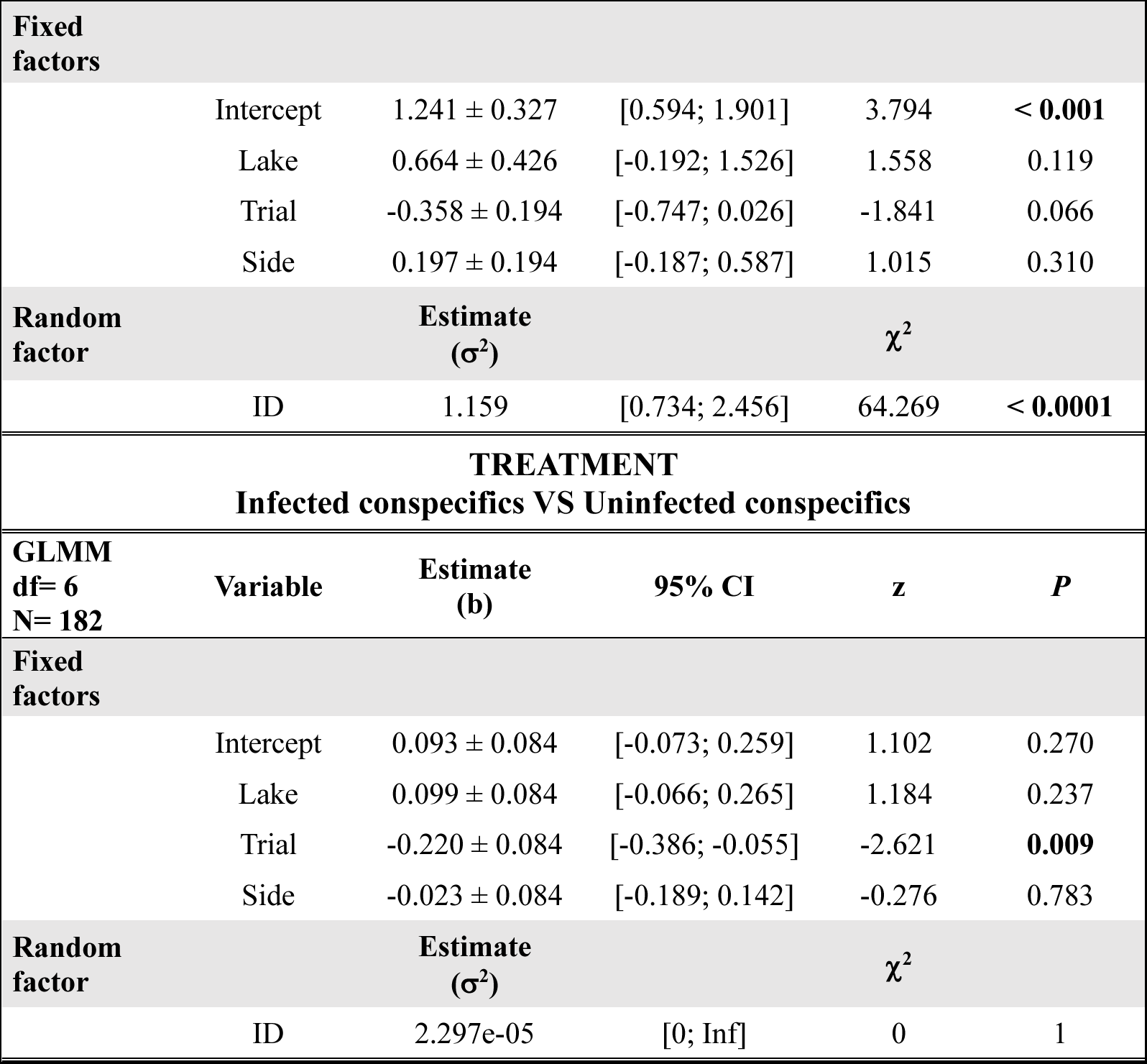
Summary of our generalized linear mixed models (GLMM) for each binary choice treatment using visual cue. Abbreviations represent the following: (df) Degrees of freedom. (N) Number of observations. (95% CI) Confidence intervals at 95%. (Lake) Lake of origin. The reference level is Lake Triton. (Trial) Trial number. The reference level is Trial 2. (Side): Side with cue. (ID) Fish identity. For the random factor, the estimate corresponds to the variance. Significant p-value are shown in bold.

**Table S3.**
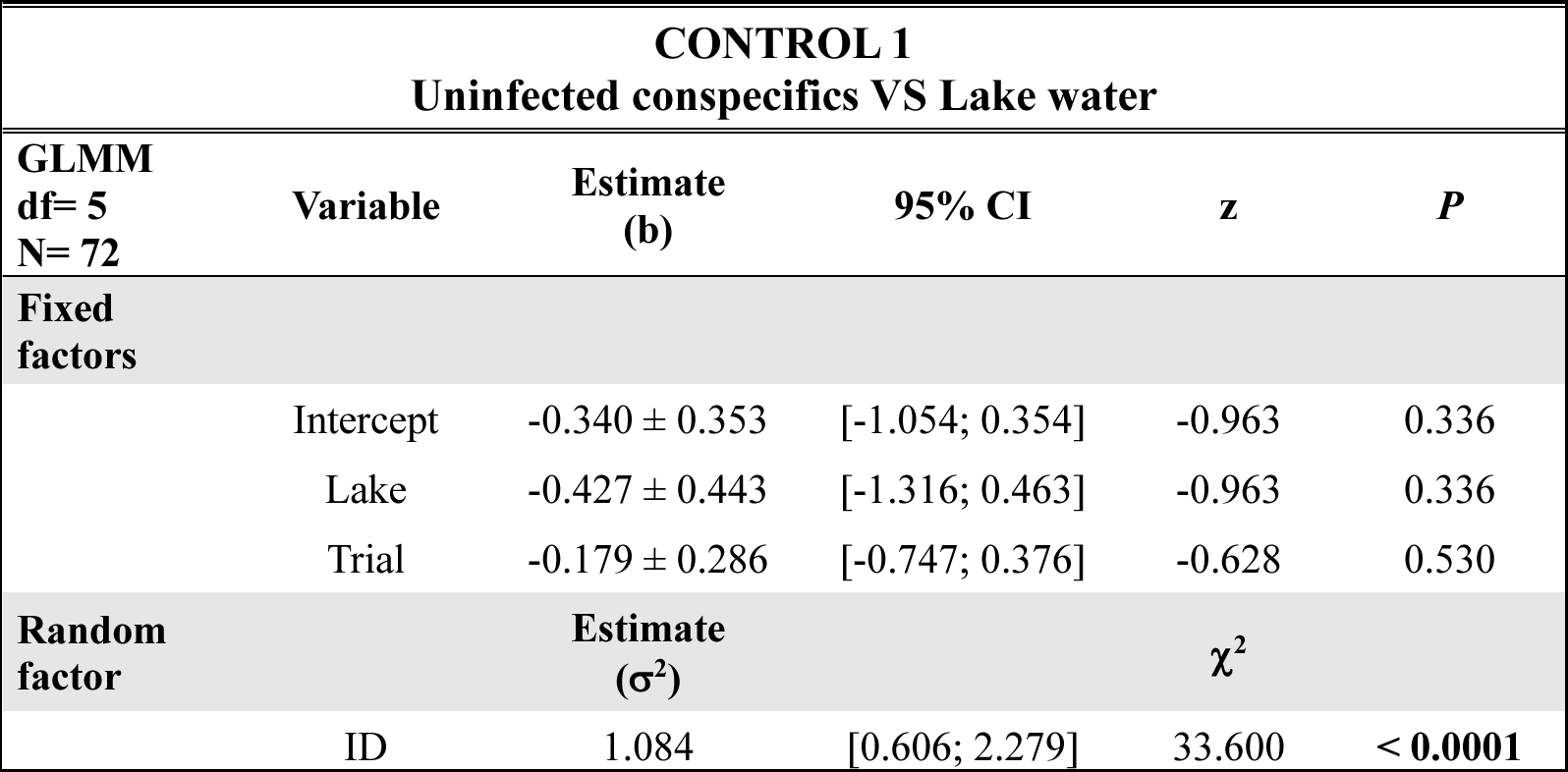
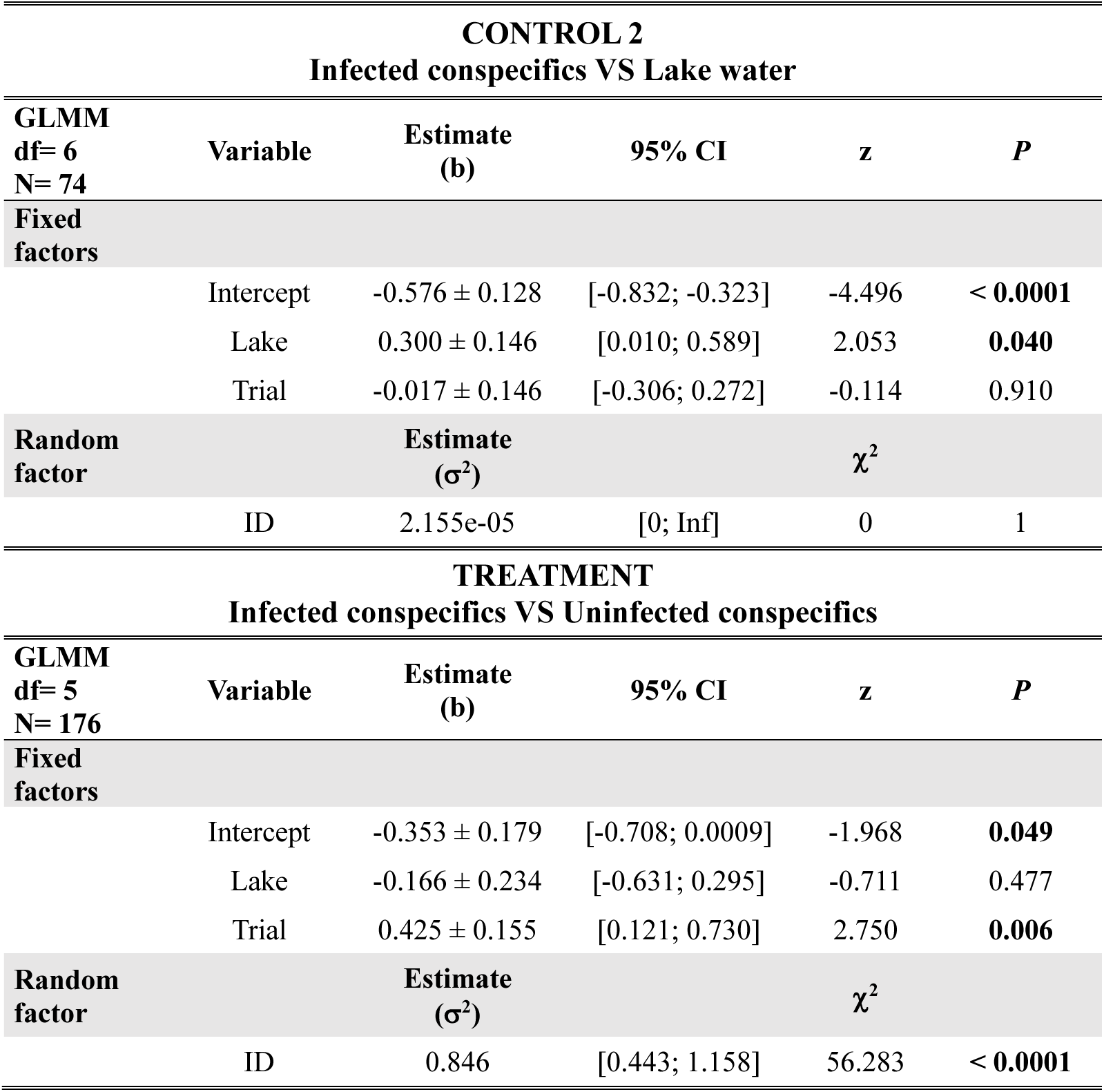
Summary of our generalized linear mixed models (GLMM) for each binary choice treatment using chemical cue. Abbreviations represent the following: (df) Degrees of freedom. (N) Number of observations. (95% CI) Confidence intervals at 95%. (Lake) Lake of origin. The reference level is Lake Triton. (Trial) Trial number. The reference level is Trial 2. (ID) Fish identity. For the random factor, the estimate corresponds to the variance. Significant p-value are shown in bold.

### Supplementary Results

To verify the impact of the trial on the preference of focal fish, we tested the preference in each of the trials individually (1 and 2) when this fixed effect was significant in our models (GLMMs). We used two-tailed paired Student’s t-tests with the proportion of time spend in each zone as the response variable (Ehman & Scott, 2001; Rahn et al., 2015, 2018). Data were tested for normality using the Shapiro-Wilk normality test and Q-Q plots on data centered and combined into a single vector. Data deviating from normality were arcsine transformed or analyzed using nonparametric paired Wilcoxon signed ranks test if still deviating from normality after being transformed.

#### Experiment 1: Visual Cues for Avoidance of Infected Conspecifics

##### Control 1: Uninfected Conspecific vs. Lake Water

**Figure S1.**
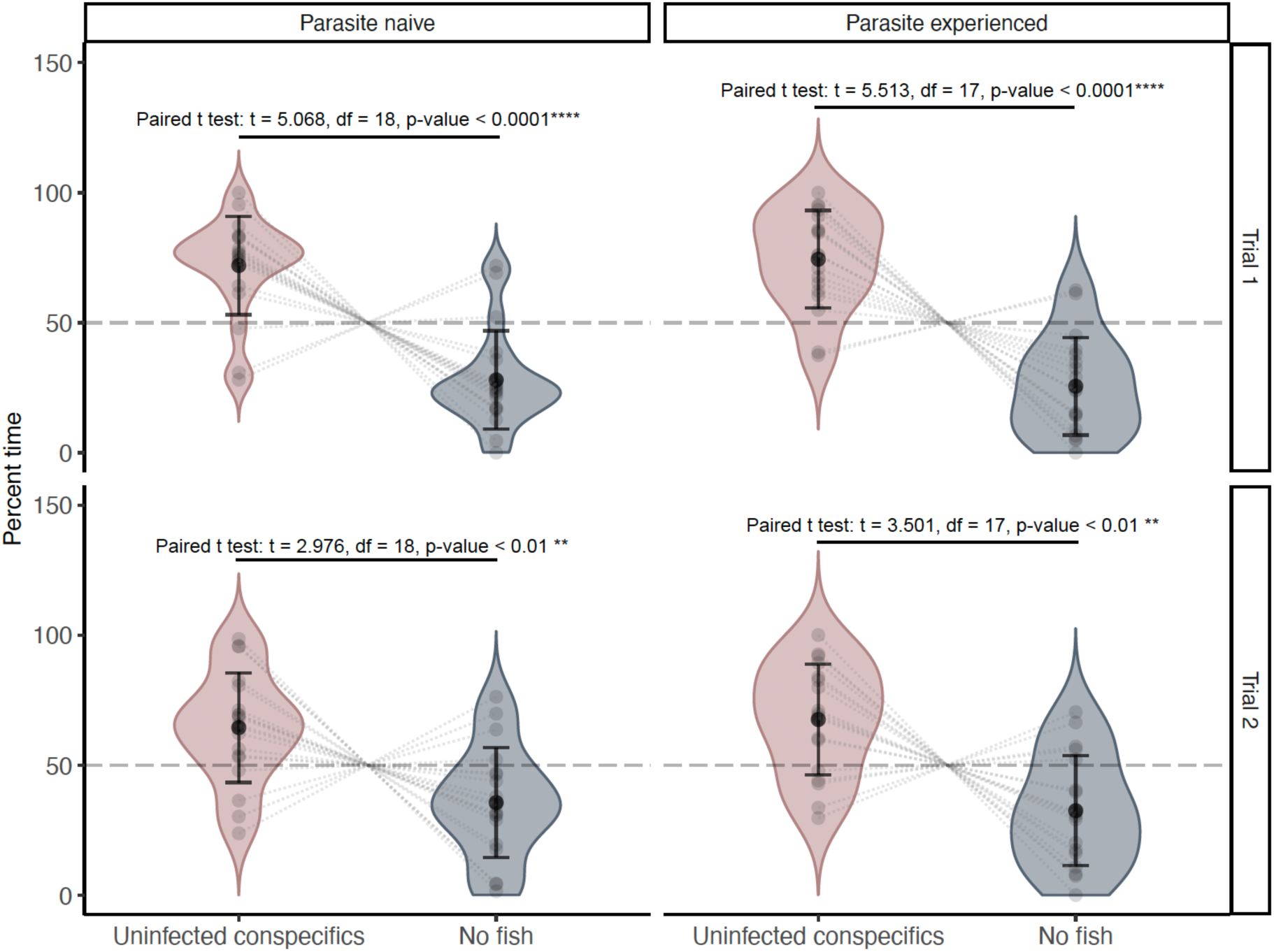
Control 1 – Visual cues divided into trials: Mean time spent with uninfected conspecifics and no fish visual cues (%) of naive (Trial 1, N = 19; Trial 2, N = 19) and experienced (Trial 1, N = 18; Trial 2, N = 18) pumpkinseed sunfish (*Lepomis gibbosus*). Dark black dots represent population means; black lines represent standard deviation; pale black dots represent observed data; two pale dots linked by a dotted line represent a single individual. Violon plots are used to show data distribution.

##### Treatment: Infected vs. Uninfected Conspecifics Cues

**Figure S2.**
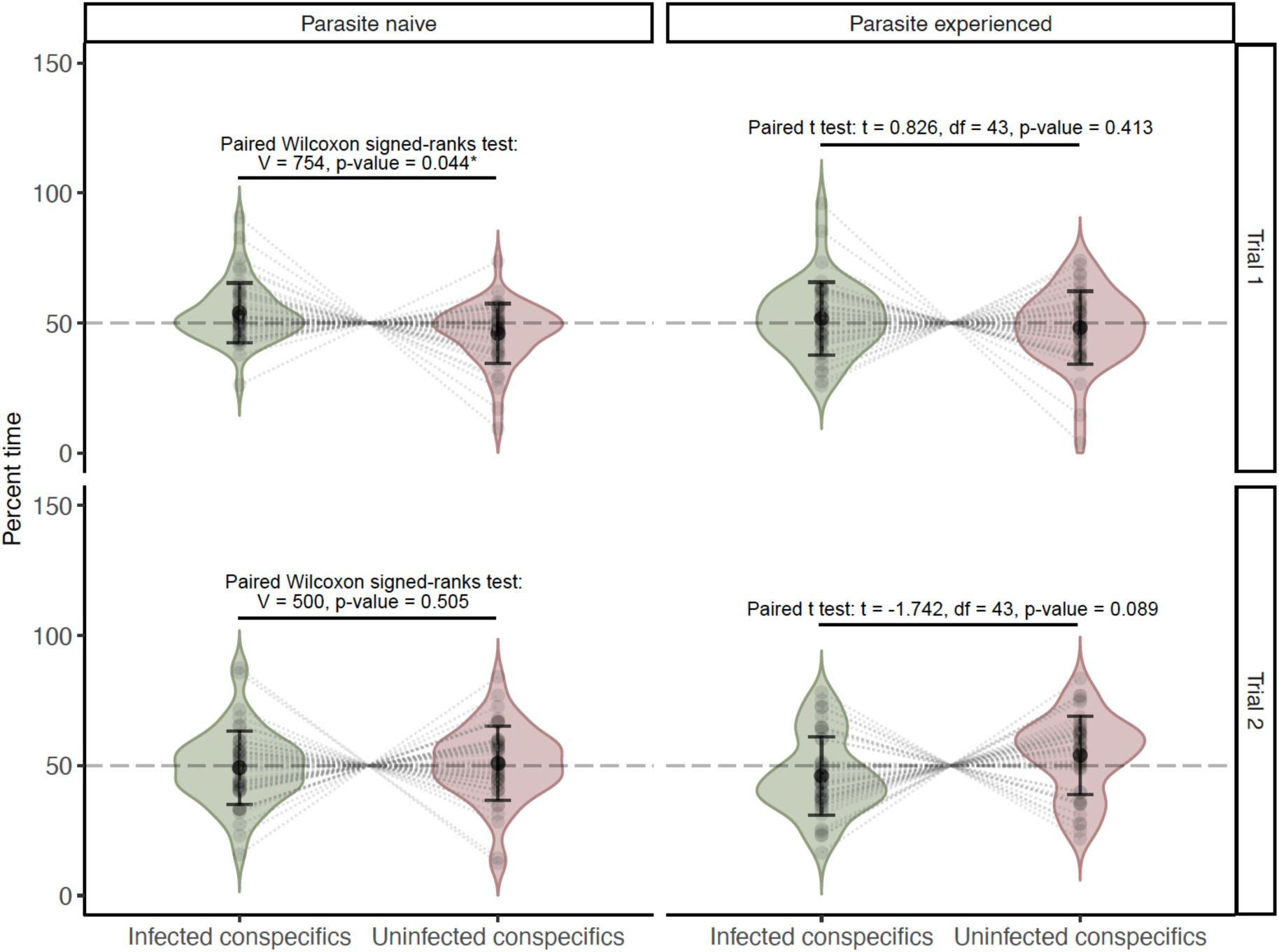
Treatment – Visual cues divided into trials: Mean time spent with infected conspecifics and uninfected conspecifics visual cues (%) of naive (Trial 1, N = 47; Trial 2, N = 47) and experienced (Trial 1, N = 44; Trial 2, N = 44) pumpkinseed sunfish (*Lepomis gibbosus*). Dark black dots represent population means; black lines represent standard deviation; pale black dots represent observed data; two pale dots linked by a dotted line represent a single individual. Violon plots are used to show data distribution.

#### Experiment 2: Chemical Cues for Avoidance of Infected Conspecifics

##### Treatment: Infected vs. Uninfected Conspecifics Cues

**Figure S3.**
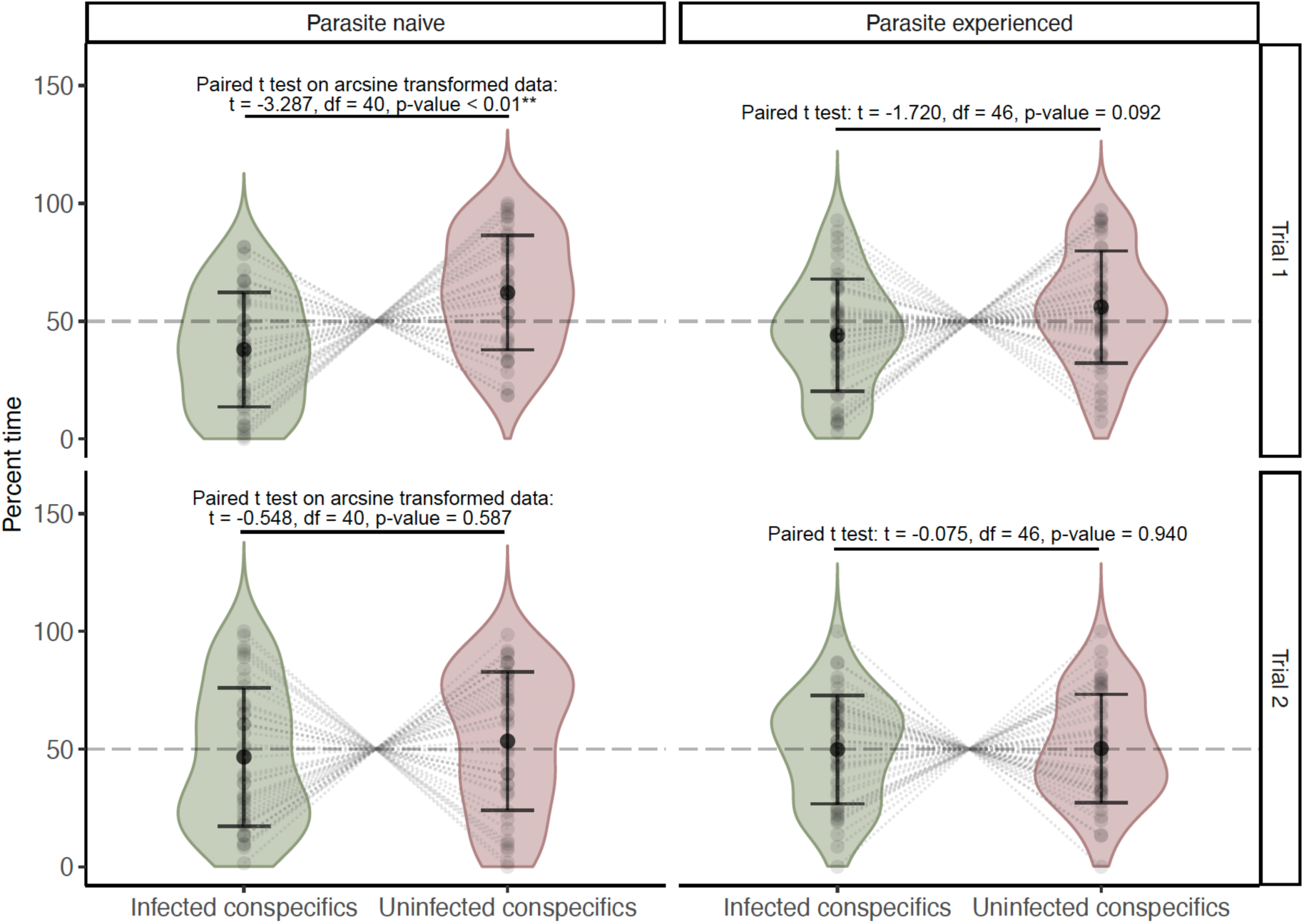
Treatment – Chemical cues divided into trials: Mean time spent with infected conspecifics and uninfected conspecifics chemical cues (%) of naive (Trial 1, N = 47; Trial 2, N = 47) and experienced (Trial 1, N = 41; Trial 2, N = 41s) pumpkinseed sunfish (*Lepomis gibbosus*). Dark black dots represent population means; black lines represent standard deviation; pale black dots represent observed data; two pale dots linked by a dotted line represent a single individual. Violon plots are used to show data distribution.

